# The Origin And Speciation Of Orchids

**DOI:** 10.1101/2023.09.10.556973

**Authors:** Oscar A. Perez-Escobar, Diego Bogarín, Natalia A.S. Przelomska, James D. Ackerman, Juan A. Balbuena, Sidonie Bellot, Roland P. Bühlmann, Betsaida Cabrera, Jose Aguilar Cano, Martha Charitonidou, Guillaume Chomicki, Mark A. Clements, Phillip Cribb, Melania Fernández, Nicola S. Flanagan, Barbara Gravendeel, Eric Hágsater, John M. Halley, Ai-Qun Hu, Carlos Jaramillo, Anna Victoria Mauad, Olivier Maurin, Robert Müntz, Ilia J. Leitch, Lan Li, Raquel Negrao, Lizbeth Oses, Charlotte Phillips, Milton Rincon, Gerardo Salazar-Chavez, Lalita Simpson, Eric Smidt, Rodolfo Solano-Gomez, Edicson Parra-Sánchez, Raymond L. Tremblay, Cassio van den Berg, Boris Stefan Villanueva, Alejandro Zuluaga, Mark W. Chase, Michael F. Fay, Fabien L. Condamine, Felix Forest, Katharina Nargar, Susanne S. Renner, William J. Baker, Alexandre Antonelli

**Author notes:** Senior authors (Oscar A. Perez-Escobar, Mark W. Chase, Michael F. Fay, Fabien L. Condamine, Felix Forest, Katharina Nargar, Susanne S. Renner, William J. Baker, Alexandre Antonelli).

## Abstract

⍰ Orchids constitute one of the most spectacular radiations of flowering plants. However, their geographical origin, historical spread across the globe, and hotspots of speciation remain uncertain due to the lack of a broad phylogenomic framework.
⍰ We present a new Orchidaceae phylogeny based on high-throughput and Sanger sequencing datasets, covering all five subfamilies, 17/22 tribes, 40/49 subtribes, 285/736 genera, and ∼7% (1,921) of the currently 29,524 accepted species. We then use it to infer geographic range evolution, diversity, and speciation patterns by adding curated geographical distribution data through the World Checklist of Vascular Plants.
⍰ Orchid’s most recent common ancestor is traced back to the Late Cretaceous in Laurasia. The modern Southeast Asian range of subfamily Apostasioideae is interpreted as relictual, matching the history of numerous clades that went extinct at higher latitudes following the global climate cooled during the Oligocene. Despite their ancient origins, modern orchid species’ diversity mainly originated over the last 5 Ma, with the fastest speciation rates found in south-eastern Central America.
⍰ Our results substantially alter our understanding of the geographic origin of orchids, previously proposed as Australian, and further pinpoint the role of Central American as a region of recent and explosive speciation.

## 1. Introduction

The angiosperm tree of life is characterised by the rise and demise of species, leading to species-rich and depauperate lineages co-existing in time and space (**Magallón *et al.*, 2019; Tietje *et al.*, 2022**). Investigating factors behind angiosperm diversification requires ancient, species-rich clades thriving across the globe and supplemented with a fossil record. One such clade is the Orchidaceae. With 29,524 species (**Chase *et al.*, 2015**; **Christenhusz & Byng, 2016; Govaerts *et al.*, 2021**), orchids are among the most species-rich groups of flowering plants, with molecular-dating studies having estimated their initial diversification at 112-76 million years ago [Ma] (**Ramírez *et al.*, 2007; Gustafsson *et al.*, 2010; Chomicki *et al.*, 2015**; **Givnish *et al.*, 2016; Serna-Sánchez *et al.*, 2021**). This diversification time is partly supported by the orchid fossil record, which includes leaf compressions and pollinaria dated from the early Eocene through the mid-Miocene found in different deposits (**Ramírez *et al.*, 2007; Conran *et al.*, 2009; Poinar, 2016a, b; Poinar & Rasmussen, 2017**). Most of these fossils have been credibly assigned to subfamilies and tribes.

An updated biogeographic study of this old, species-rich, and cosmopolitan family (**Fig. 1a-d)**, which is uniquely diverse in pollination systems (**Ackerman *et al.*, 2023**) and adaptations to different habitats (Fig. **1e-f**), is an essential step towards a better understanding of monocot clades with wide distribution ranges and diversity of adaptations (e.g., Arecaceae [**Couvreur *et al.*, 2011**], Bromeliaceae [**Benzing, 2000**], Poaceae [**Soreng *et al.*, 2015**]). Such a study can build on several in-depth analyses that have focused on subclades (**Bouetard *et al.*, 2010**; **Guo *et al.*, 2012; Freudenstein & Chase, 2015; Pérez-Escobar *et al.*, 2017a; Nauheimer *et al.*, 2018**). In a benchmark study of the biogeographic history and diversification of the entire family, **Givnish *et al.* (2016**) used a fossil-calibrated plastid tree for 173 genera (out of 736 in total) of orchids representing all five subfamilies, along with selected outgroups in the Asparagales (Asteliaceae, Blandfordiaceae, Boryaceae, Hypoxidaceae, Iridaceae, Lanariaceae). Using a likelihood approach (**Matzke, 2013**) and a single consensus ultrametric tree, their biogeographic analyses relied on a geographic matrix where terminals, usually representing genera, were coded for the entire distribution of the respective genera. Their results pointed to an origin and initial diversification of orchids in Australia during the mid-Cretaceous, 120–90 Ma. Notably, their range estimates for the Orchidaceae stem node indicated Australia as ∼40% likely, Neotropics plus Australia as ∼20%, and all other ranges together as ∼40%. At 90 Ma, Australia was at high latitudes connected to Antarctica and part of the southern supercontinent, Gondwana, with connections to South America, and the authors, therefore, stressed the likely importance of expansion across Antarctica. A Gondwanan origin for Orchidaceae was also suggested by **Chase (2001)**. The extent to which taxon sampling, phylogenetic uncertainty, and the chosen approach for coding geographical ranges influenced the estimation of ancestral areas in Givnish *et al.*’s study remains unknown.

**Figure 1.**
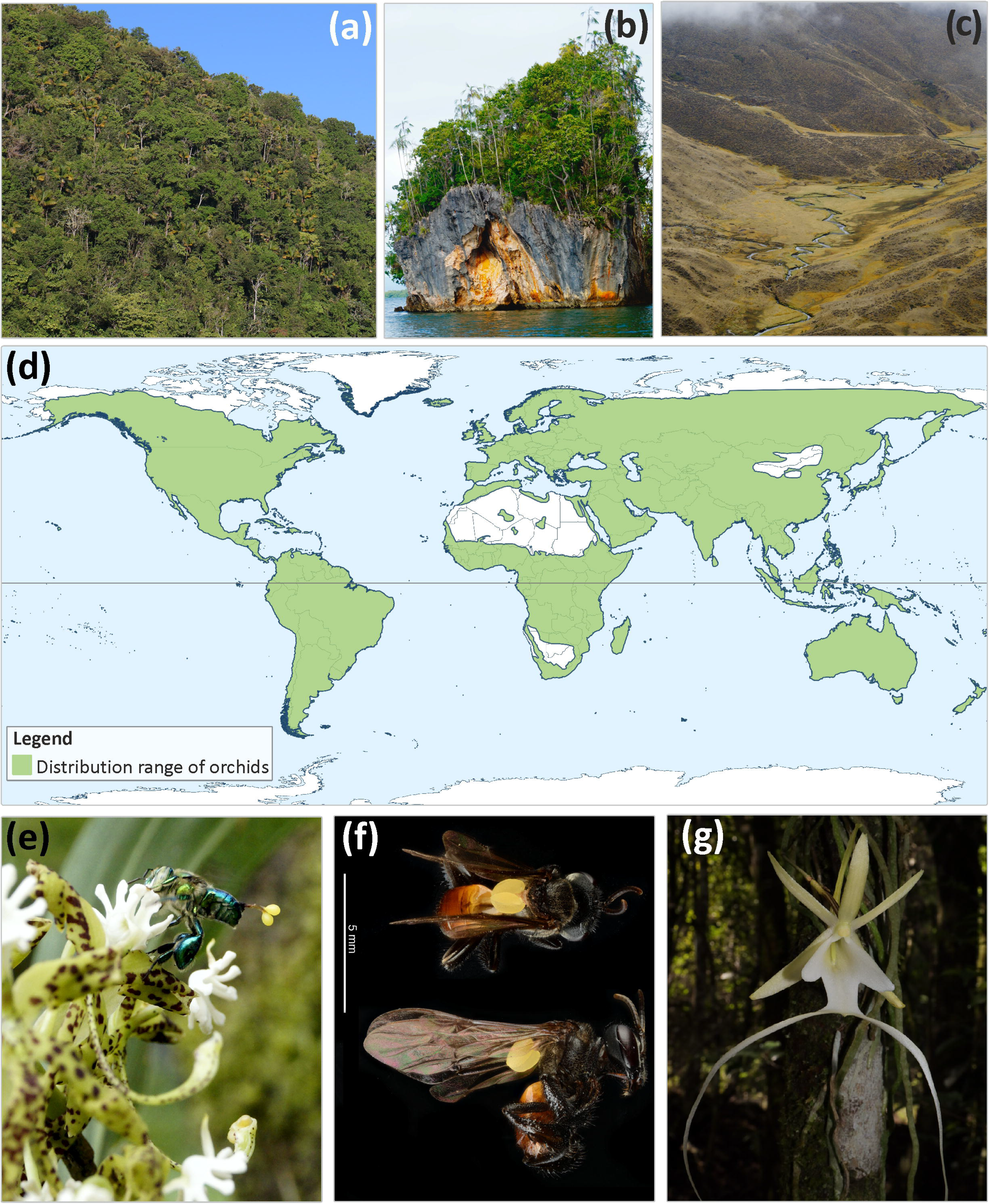
Diversity of orchid habitats (A-C), modern distribution range (D) and pollination systems in Orchidaceae (E-G). **A**) Tropical cloud forests; **B**) Tropical lowland wet forests; **C**) High-elevation grasslands; **E**) the Neotropical epiphyte *Cycnoches guttulatum* Schltr., pollinated by male euglossine bees collecting aromatic compounds; **F**) *Trigona* bee carrying a pollinarium of a Neotropical *Xylobium elongatum* (top: dorsal view; bottom: side view); G) the epiphytic and leafless *Dendrophylax sallei* (Rchb.f.) Benth. ex Rolfe., a Neotropical orchid pollinated by moths. Photos: Oscar A. Pérez-Escobar & Diego Bogarín.

**Givnish *et al.* (2016: Table 1)** also addressed the correlates of diversification rates through time and inferred Southeast Asia as the region with the highest net diversification rates. Studying how diversification rates are linked to geographical variation helps understand not only the pace at which extant species diversity has accumulated but also the correlated biotic and abiotic variables (**Condamine *et al.*, 2013**; **Velasco & Pinto-Ledezma, 2022**). Although orchid species diversity is clearly unevenly distributed across their distribution (**Vitt *et al.*, 2023**), no study has yet assessed the relationship between the distribution of orchid species richness and their underlying current speciation rates at a global scale for the entire family.

Here, we revisit the biogeographic history of Orchidaceae and infer geographic patterns of speciation, using a greatly expanded taxon sampling (1,921 species, 285 genera) as compared to previous phylogenomic studies, with a particular focus on early divergences. We generate a robust phylogenomic framework by combining high-throughput and Sanger sequencing data. By employing multiple topologies as input for the phylogenetic comparative analyses, we account for the effects of branch length variation and uncertainty in the position of weakly supported lineages. Our spatial information derives from herbarium-vouchered, georeferenced occurrence data, sourced from the Global Biodiversity Information Facility (GBIF; https://www.gbif.org) and the RAINBIO mega-database (**Dauby *et al*., 2016**). This dataset was subsequently vetted using the spatial distribution data of the World Checklist of Vascular Plants (WCVP; **Govaerts *et al*., 2021**). Based on these datasets, we: *i*) infer the evolutionary and biogeographical history of the orchid family, *ii*) test the hypothesis of an Australasian origin for the most recent common ancestor of living orchids, and *iii*) revisit the hypothesis that Southeast Asia is the region with the highest current diversification of orchids.

## 2. Material and Methods

### 2.1 Taxon sampling, DNA library preparation and sequencing

Our phylogenomic framework was assembled using a high-throughput-sequencing dataset, complemented with Sanger data for single genes. The high-throughput-sequencing dataset focused on the genes targeted by the Angiosperms353 probe set (**Johnson *et al.*, 2019; Baker *et al.*, 2022**) and includes 448 orchid species, representing 285/736 genera, 40/49 subtribes, 17/22 tribes, and 5/5 subfamilies (in the classification of **Chase *et al.*, 2015**). These data were generated by the Plant and Fungal Trees of Life Project (**Baker *et al.*, 2022**) at the Royal Botanic Gardens, Kew, and the Genomics for Australian Plants Consortium (https://www.genomicsforaustralianplants.com/). Nineteen monocot species were included as outgroup taxa, with *Dioscorea caucasica* (Dioscoreales) used as a rooting terminal, for a total of 467 species in the high-throughput-sequencing dataset. Illumina sequencing reads were newly produced for 377 samples (*vs* **Pérez-Escobar *et al.*, 2021**) from expertly curated specimens housed in the Kew (K) and Australian National herbaria (CANB), including 23 species sampled from type material and silica-gel dried tissue (**Table S1**). Detailed protocols for DNA extraction and total genomic DNA libraries are provided in **Methods S1**.

The Sanger sequencing dataset was assembled through the SuperCRUNCH pipeline (**Portik *et al.*, 2020**). This pipeline was executed using an initial set of 24,172 sequences of the nuclear ribosomal ITS and the plastid *mat*K markers – two commonly sequenced, informative loci for Orchidaceae and other angiosperms (**Grace *et al.*, 2021**) – obtained from GenBank (https://www.ncbi.nlm.nih.gov/nuccore). The following parameters were used: *i*) search terms of ‘ITS; ITS1; ITS2; internal transcribed spacer 1; partial sequence; 5.8S ribosomal RNA gene; complete sequence; internal transcribed spacer 2’ for ITS, and ‘*matK*; *trnK*; maturase K; *trnK* gene, intron; and maturase K *matK* gene’ for *matK*; *ii*) a list of 285 generic (accepted) names that were sampled in the high-throughput-sequencing dataset; and *iii*) similarity searches between sequences using MEGABLAST (**Camacho **et al.*,* 2009**), retaining a maximum of 200 hits per search. After removing duplicated sequences and obvious contaminants (defined as sequences with misplaced positions given current taxonomic views), we retained sequences of nrITS and plastid *matK* for 2,060 species, of which 64% had sequences produced from the same voucher specimen (**Table S2**).

### 2.1 High-throughput and Sanger sequencing data analyses

Illumina libraries were quality-assessed with FastQC software v.0.11 (available at https://www.bioinformatics.babraham.ac.uk/projects/fastqc/). Paired-end reads were quality-filtered and adapter-trimmed using TrimGalore! V.0.6.4 available at (https://github.com/FelixKrueger/TrimGalore), using the following parameters: *i*) *-q 30* (minimum Phred score), *ii*) *–length* 20 (minimum read length), *iii*) retaining read pairs that passed the quality thresholds. In-silico retrieval of the Angiosperms353 coding loci was conducted using the pipeline HybPiper v.2.1.6 (**Johnson **et al.*,* 2016**) and the same parameters and software described in **Pérez-Escobar *et al.* (2021a;** also see *Methods S1*). Next, low-copy nuclear and Sanger markers were aligned using the software MAFFT v.7.4 (**Katoh & Standley, 2013**) in conjunction with the iterative refinement method FFT-NS-I, implementing a maximum of 1,000 iterations. To account for potential biases from missing data in phylogenetic analyses, sequences shorter than 50% of the total alignment length were excluded. Subsequently, alignments were filtered for misaligned positions using TAPER v.1.0 (**Zhang **et al.*,* 2021**), using a cut-off value of 1 (flag *–c*), and subsequently inspected by eye with Geneious v.8.0 (available at https://www.geneious.com/). The Sanger/Angiosperms353 nucleotide alignments are accessible at (at10.6084/m9.figshare.22245940).

### 2.2 Distance-based, maximum likelihood, Bayesian, and multispecies coalescence phylogenomic inferences

We computed maximum likelihood (ML) trees from the individual Sanger and Angiosperms353 nucleotide alignments using RAxML v.8.0 (**Stamatakis 2014**) with the following parameters: *i*) 500 rapid bootstrap replicates (flags *-#* 500 and *-x*); *ii*) the GTR+ Γ nucleotide substitution model. Additionally, a multispecies coalescent (MSC) tree was inferred from the individual Angiosperms353 gene trees produced by RAxML using the software ASTRAL-III v.5.6 (**Zhang **et al.*,* 2018**) and the flag –t 2 (full tree annotation), after collapsing bipartitions with a likelihood bootstrap percentage (LBP) < 20% (achieved with the function new_ed of the package *newick_utils* v.1.6.0, available at https://bioweb.pasteur.fr/packages/pack@newick-utils@1.6). To visualise the proportion of gene-tree quartets that agreed with the species tree, we produced quartet support pie charts for every branch represented in the species’ tree, using the full annotation produced by ASTRAL-III. We also visualized the proportion of gene trees in agreement with each species’ tree bipartition using Phyparts v.1.0, by labelling as informative any gene-tree bipartition with a LBP > 20% (**Smith **et al.*,* 2015**).

Following the overall topological congruence amongst the ITS and *mat*K datasets and Angiosperms353 low-copy nuclear gene alignments revealed by the MSC and a goodness-of-fit test (**Balbuena **et al.*,* 2013**; **Pérez-Escobar **et al.*,* 2015**, see *Methods S1* and *Results and Discussion S1*), we also computed ML phylogenetic trees from *a*) a super-matrix derived by concatenating the ITS and *matK* sequences, *b*) a super-matrix derived from the concatenation of the Angiosperms353 loci using RaxML v.8.0 and the same settings previously specified, considering each super-matrix as a single partition. Lastly, we inferred a consensus network using the 500 bootstrap replicates produced from the Angiosperms353 super-matrix by RAxML v8.0, using SplitsTree v.4.0 (**Huson & Bryant, 2006**), a mean edge weight, and a threshold value of 0.75 to filter out any split absent in such a proportion of the bootstrap trees. The alignments, trees, and consensus network are accessible at (at10.6084/m9.figshare.22245940).

### 2.3 Molecular clock dating analyses and species-level phylogeny assembly

Absolute age estimation analysis of orchid phylogenies was conducted in two stages, and by using a Bayesian framework implemented in BEAST v.2.6 (**Bouckaert **et al.*,* 2019**), as follows:

***(i)*** The backbone was inferred by subsampling the Angiosperms353 low copy nuclear gene alignments using SortaDate v.1.0 (**Smith **et al.*,* 2018**). Here, we selected the top 25 low-copy nuclear gene alignments with the lowest root-to-tip variance coefficient (i.e., clock-likeness). This selection ensured the representation of the entire generic diversity as sampled by the ML high-throughput dataset (i.e., 285 genera; see *Methods S1*, and **Table S3, S4**). This data subset was imported in BEAUTi v.2.6 (**Bouckaert **et al.*,* 2019**) as unlinked partitions, with the same priors employed by Pérez-Escobar *et al.* (**2021b**), as follows: *a*) the GTR nucleotide substitution model and a rate heterogeneity among sites modelled by a Γ distribution with four categories, *b*) an uncorrelated log-normal relaxed molecular clock in combination with a prior clock rate interval of 0.0001-0.001 substitutions/site/Ma, modelled by an uniform distribution, *c*) a birth-death tree process, modelled by a uniform distribution for the birth and relative death rates, *d*) three secondary calibration points retrieved from Givnish *et al.* (**2015**), modelled by normal distributions (σ = 1) and located at the root node, i.e., most recent common ancestor (MRCA) of Dioscoreales, Orchidaceae and Liliales (125 Ma), the MRCA of Orchidaceae (89 Ma) and the MRCA of subtribe Goodyerinae (32 Ma), *e*) the ML consensus phylogram of generic lineages produced in RaxML v.8.0 as a starting tree, *f*) 500 million generations, sampling every 100,000 generations and ensuring that all posterior values reached effective sample sizes (ESS) > 100. The proportion of informative and missing sequences per taxon sample is provided in **Table S4.** Finally, the support for the backbone ultrametric trees derived from BEAST was assessed by first computing a maximum credibility consensus tree from the Markov chain Monte Carlo (MCMC) posterior trees using TreeAnnotator v.2.6 (https://www.beast2.org/treeannotator/), using a burn-in threshold of 10%, and then counting the proportion of bipartitions falling on different support intervals.
***(ii)*** The species-level-ultrametric phylogeny of Orchidaceae was produced by inputting the ITS and *matK* alignments as unlinked partitions in BEAUTi v.2.6 (**Bouckaert **et al.*,* 2019**), using the same priors described in (*i*). Furthermore, the ML consensus phylogram produced in RaxML v.8.0 from the ITS-*matK* supermatrix was used as a starting tree.
***(iii)*** To account for phylogenetic uncertainty, we relied on constructing multiple ultrametric species trees through a novel pipeline (**Fig. 2**) as opposed to using a single consensus maximum clade credibility tree. We first randomly sampled 10 posterior trees derived from the BEAST analyses conducted on the genus-level Angiosperms353 and the ITS-*mat*K Sanger species-level datasets. Then, for each genus represented in the Angiosperms353 chronograms, we pruned its counterpart clade sampled on the species-level Sanger chronogram and proceeded to graft it onto the corresponding stem of the Angiosperms353 chronogram. This operation was conducted on each pair of randomly sampled posterior trees (hence called PP species trees). A detailed description of the pipeline is provided in the supplementary *Methods S1*.

**Figure 2.**
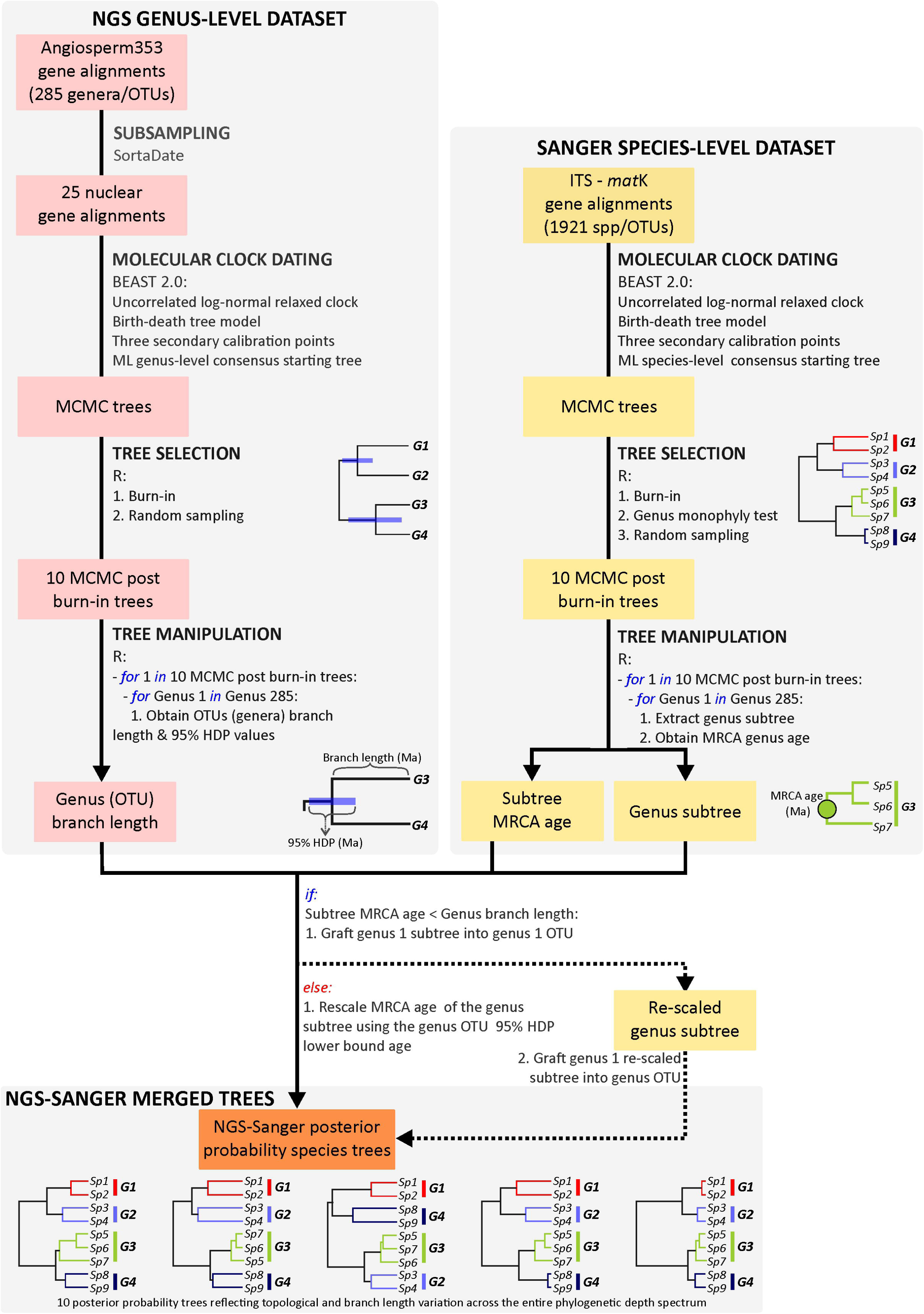
An overview of the pipeline used to produce ultrametric NGS-Sanger merged posterior probability species trees (PP species trees). The pipeline uses as input a custom set of randomly sampled MCMC trees derived from independently conducted molecular clock dating analyses on the genus-level Angiosperms353 and species-level Sanger gene alignments. It then produces a custom set of species-level trees that reflect branch length variation and support in deep (tribe, subtribe and genus) and recent (species) time as informed by the Angiosperms353 and Sanger datasets. A detailed description of the pipeline is provided in the supplementary *Methods S1*.

### 2.4 Biogeographic analyses

Biogeographical state estimations were conducted on the 10 PP species trees. These were produced by pruning and grafting the Sanger ultrametric species trees onto the Angiosperms353 genus-level chronograms. We estimated areas of origin and geographic range using the ML approach of dispersal-extinction-cladogenesis (DEC, **Ree & Smith, 2008**) as implemented in the C++ version (**Beeravolu & Condamine, 2016**). The eight geographic areas considered were: (1) Palearctic, defined as Europe, Siberia, Central Asia, and Western Asia; (2) Nearctic, including all North America to the north of Tehuantepec in Mexico and excluding the southern tip of Florida; (3) Neotropics, including Central America, the Caribbean Islands, and South America; (4) Africa, defined as the whole African continent including Madagascar, and Arabian Peninsula; (5) Indomalaya, including India, southern China, and Wallacea; and (6) Australasia, defined as everything east of Heilprin-Lydekker’s Line (**Ali & Heaney, 2021**). A time-stratified geographic model specified constraints on area connectivity by coding 0 if any two areas are not connected or 1 if they are connected at a given period based on paleogeographic reconstructions. The first time-slice covers the Late Cretaceous and early Paleocene (100.5-60 Ma), corresponding to a time where Gondwana and Laurasia were already separated (**de Lamotte **et al.*,* 2015**). The second time-slice covers the late Paleocene to early Oligocene (60–30 Ma), and the third covers the early Oligocene to the present (30-0 Ma). To avoid bias results supporting ancestral areas in favour of hyperdiverse biomes, our sampling per biogeographic area used proportional sampling that reflected known diversity patterns across biogeographic realms, with the Asian and American tropics hosting the highest levels of species richness (**Vitt **et al.*,* 2023**). Here, we included 905 species distributed in the Neotropics (6% of total known diversity in the area; **Govaerts **et al.*,* 2021**), 623 from the Indomalaya region (6%), 347 from the Palearctic (5%), 221 from the Afrotropics (9%), 180 from Australasia (15%), and 44 from the Nearctic (10%).

### 2.5 Spatial analysis of species diversity and speciation rates

We queried 29,524 accepted orchid species names obtained from the WCVP through the GBIF (https://www.gbif.org/) and RAINBIO (https://gdauby.github.io/rainbio/index.html; **Dauby **et al.*,* 2016**) databases. Our initial step involved accessing the GBIF database through the R-package SPOCC (available at https://github.com/ropensci/spocc). Using the *occ* function, we downloaded up to 1,000 records for each queried species name, ensuring that we only selected records linked to geographical coordinates and preserved specimens. The RAINBIO repository was manually accessed. Next, we conducted an automated filtering using the SpeciesGeoCodeR package (**Töpel **et al.*,* 2017**) to exclude duplicated records and those located within urban areas. Subsequently, we removed occurrences that fell outside global coastlines. Lastly, to mitigate the influence of misidentified records on downstream analyses (**Maldonado **et al.*,* 2015**), we further filtered out records for which the distribution did not match with the species distribution per botanical country (level 3 of the World Geographical Scheme for Recording Plant Distributions; **Brummitt, 2001**) provided by the WCVP. The original record database sourced from GBIF is available at https://doi.org/10.15468/dl.v2gwxv, and the filtered geographical records used in all downstream analyses will be available at (*forthcoming FigShare link*).

Estimating net diversification rates (speciation [λ] minus extinction [*µ*]) in plant lineages remains challenging (**Louca & Pennell, 2021**). However, it is possible to estimate λ from a densely sampled and robust phylogenetic tree, if the provided sampling fraction included in that tree is known, and the phylogenetic framework is extensive (**Louca & Pennell, 2021**). Our approach focuses on speciation dynamics, as inferred from tip λ rates. This metric represents contemporary rates of speciation for a given lineage and is less prone to bias than net diversification rates (**Title & Rabosky, 2019**). To obtain tip λ rates, we fitted a time-dependent model using BAMM v.2.5.0 (**Rabosky **et al.*,* 2013**). This was informed by sampling fractions, contrasting accepted species number per genus from the WCVP database with the number of species sampled in our phylogenetic analyses. Here, 54% of the genera sampled attained a species representativeness of 10-50%, with 21% genera including sequences of > 50% of the known species diversity, and 25% sequences representing < 10% of the known species diversity. This analysis was conducted on each of the 10 PP species trees generated by pruning and grafting the genus- and species-level chronograms produced by BEAST. We initially computed prior values for the initial λ and shift parameters, using the R package BAMMtools (**Rabosky **et al.*,* 2014),** and performed 10 million generations, sampling the MCMC simulations every 10,000 generations. The BAMM output was analysed in BAMMtools to ensure that all analyses reached convergence (ESS values > 100). Mean speciation rates at present time (λ tip rates) were extracted for all terminals of the phylogenetic trees using the *getTipRates* function in BAMMtools, and then linked to the filtered geographical species occurrences.

Global patterns of species richness were inferred by calculating the average number of species per political and botanical country, and per grid cell (100 km × 100 km), using the filtered occurrence records, as well as the average and maximum λ rate values per grid cell. All calculations were performed in the open source software QGIS v.3.0 (available at https://www.qgis.org/en/site/forusers/download.html) and the R package KEWR (Walker 2022).

## 3. Results and Discussion

### 3.1. A new phylogenomic framework for the orchid family

This study underlines the advantage of combining high-throughput and Sanger sequencing datasets to achieve denser taxon sampling at relatively lower costs, while accounting for phylogenetic uncertainty. Our MSC and ML analyses yielded a well-supported phylogenomic framework for the Orchidaceae, containing 38% of the currently accepted orchid genera (Fig. **3**; **Fig. S1-S8; see Results and Discussion S1**). The relationships among subfamilies and tribes largely agree with findings from a recent study by **Zhang *et al.* (2023)**. This study presented a phylogeny of the Orchidaceae produced from 610 to 1,195 nuclear transcriptomes obtained from 437 species representing 131 genera. Notable exceptions include the monophyly of Bletiinae within Epidendreae, and the position of Eriopsidinae within the Cymbidieae (Figure S2 in **Zhang **et al.*,* 2023**). A detailed view of the summary trees produced by **Zhang *et al.*** reveals evidence of intragenomic conflicts (see Figure S3): 46 bipartitions in the MSC trees attained low local posterior probabilities (< 0.5), and LBPs (<70%), suggesting that dominant alternative relationships exist for the branches in question. Additionally, **Zhang *et al.*** recovered the subtribe Bletiinae as paraphyletic (i.e., *Bletia* and *Chysis* as successive sister to Ponerinae, Laeliinae and Pleurothallidinae), a relationship recovered with low support by **van den Berg *et al.* (2005)** and **Górniak *et al.* (2010**). Unfortunately, voucher information for the data of **Zhang *et al.* (2023)** has not been made publicly available (at the time of writing), which limits the extent to which these phylogenomic discrepancies can be evaluated (PRJNA923320 accession code, queried on 25 August 2023 in https://www.ncbi.nlm.nih.gov/, returned zero results).

**Figure 3.**
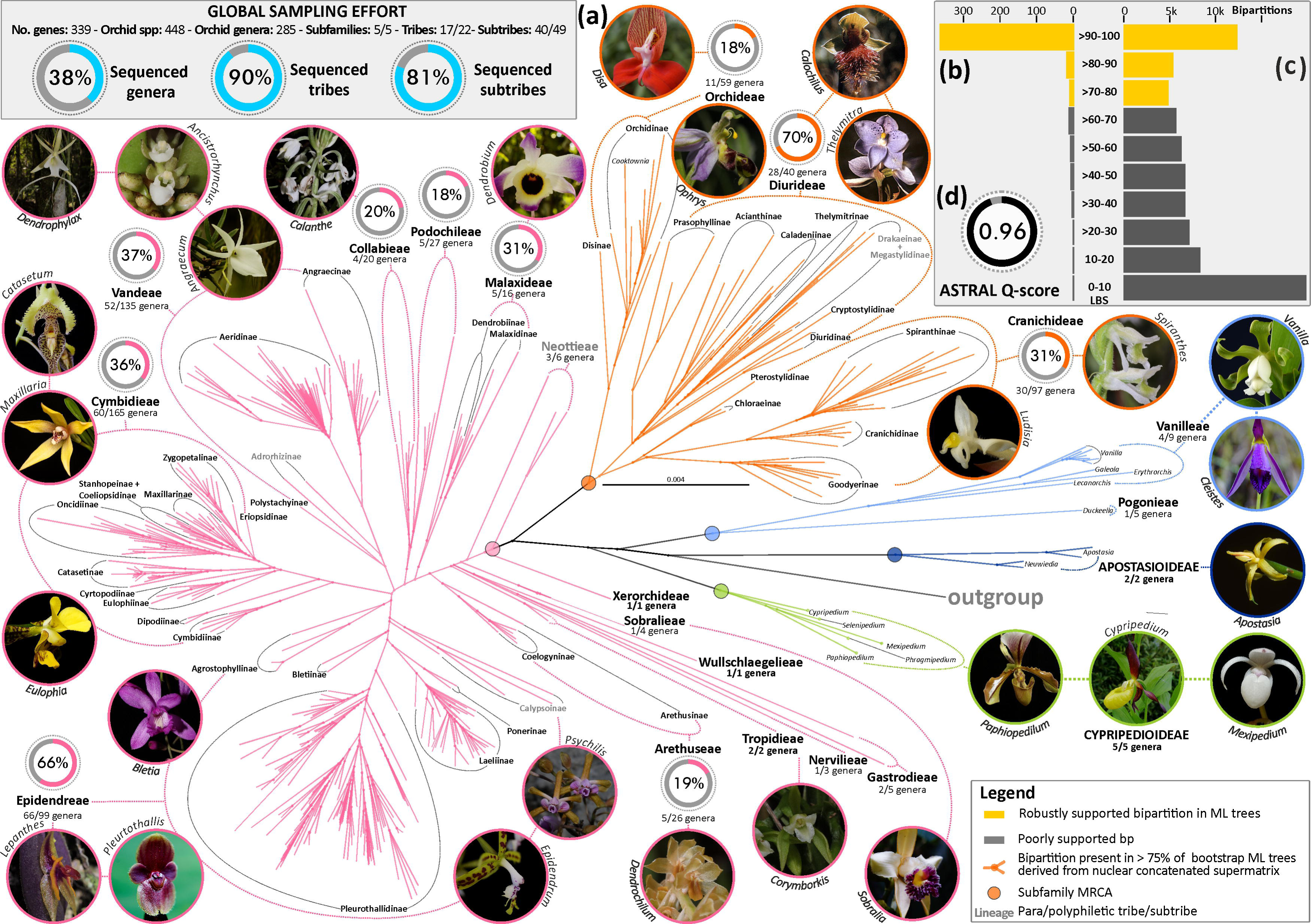
Phylogenetic relationships of Orchidaceae. **A**) Consensus tree-network inferred from 200 bootstrap replicate maximum likelihood (ML) trees derived from the concatenated alignment of 339 low-copy nuclear genes. Circles at nodes represent bipartitions present in > 75% of the bootstrap ML trees (see Figures S1-S6 for a detailed on the consensus tree-network). Non monophyletic groups are highlighted in bold and grey. Samples sequenced from typological material are highlighted in bold and pink. **B**) Number of bipartitions (bp) with different likelihood bootstrap support (LBS) values derived from an ML tree inferred from the 339 low-copy nuclear gene alignment. **C**) Number of bp with different LBS values derived from 339 nuclear ML gene trees. Grey bars represent poorly supported bipartitions, whereas yellow bars represent moderately to maximally supported bipartitions. **D**) Normalised Q-score value derived from an ASTRAL-III analysis, computed from 339 nuclear ML gene trees. Photos: Oscar A. Pérez-Escobar, Diego Bogarín, Sebastian Vieria, Kerry Dressler.

Our results unambiguously place in the orchid tree of life the monospecific genus *Cooktownia* D.L.Jones (Orchidoideae) and the small genus *Claderia* (two species; Epidendroideae). These genera are distributed from the western Indian Ocean to the western Pacific and from the south-western Pacific to New Zealand. Previous classifications placed *Cooktownia robertsii* in the tribe Orchideae, based on stigmatic chamber characters (**Jones 1997**). Our findings suggest that *Cooktownia* is most closely related to *Habenaria* Willd., a large, almost cosmopolitan genus from subtribe Orchidinae. On the other hand, *Claderia* is confirmed as sister to *Agrostophyllum* Blume and *Earina* Lindl. (Agrostophyllinae, Epidendreae). We obtained sequences from the type specimen of *Claderia viridiflora* collected in 1867. Though the Angiosperms353 gene recovery was predictably low for this specimen, we successfully identified 40 genes, with 23 being informative (**Table S1**). *Claderia viridiflora* had been included in the predominantly African subtribe Eulophiinae based on morphological similarities in the gynostemium and unpublished nrITS sequences (**Cribb & Pridgeon, 2009**). Our placement in Agrostophyllinae agrees with **Niissalo **et al.*,* (2023),** who recovered *Claderia* as a sister to *Agrostophyllum* based on plastid genomes and the nrITS loci, confirming the robustness of our findings.

### 3.2. A Late Cretaceous, Laurasian origin of Orchidaceae and the relictual range of apostasioids

The 10 PP species trees robustly supported relationships and revealed no major conflicts with the MSC tree produced in ASTRAL, with the sole exception of the position of *Corymborkis*, which, in some instances, was recovered as nested within tribe Gastrodieae (**Fig. S7, S8**). The stem age of Orchidaceae is estimated to be around 120 ± 6 Ma (Fig. **S7**) and the crown age around 83 ± 10 Ma, corresponding to the Early and Late Cretaceous, respectively, in agreement with previous estimates (**Chase, 2001; Ramírez **et al.*,* 2007**; **Chomicki **et al.*,* 2015**; **Givnish **et al.*,* 2016; Zhang **et al.*,* 2023**). However, our results indicate that the majority of present-day orchid species diversity originated over the past 5 million years (Fig. **S9**).

Seven of the biogeographical estimates conducted on the PP species trees supported Laurasia (Nearctic + Palearctic) as the place of the initial diversification of extant orchids (relative probability = 0.15-0.30; **Table S5**), equivalent to their most recent common ancestor (Fig. **4a**; Table S5). Conversely, the three other estimates placed the orchid MRCA in Laurasia + Neotropics or in Gondwana (Neotropics + Australasia + Antarctica) (relative probability = 0.14-0.19). The second most likely areas had nearly equal probabilities to those for the first most likely area (relative probability = 0.14-0.18), all placing the orchid MRCA in Laurasia. This matches the important role that Laurasia played during the late Cretaceous, fostering the initial diversification of diverse tropical and warm temperate flowering plant lineages. Notable examples include the palm family Arecaeae (**Baker & Couvreur, 2013**) and the yams, *Dioscorea* (Dioscoreaceae: **Viruel **et al.*,* 2016**).

**Figure 4.**
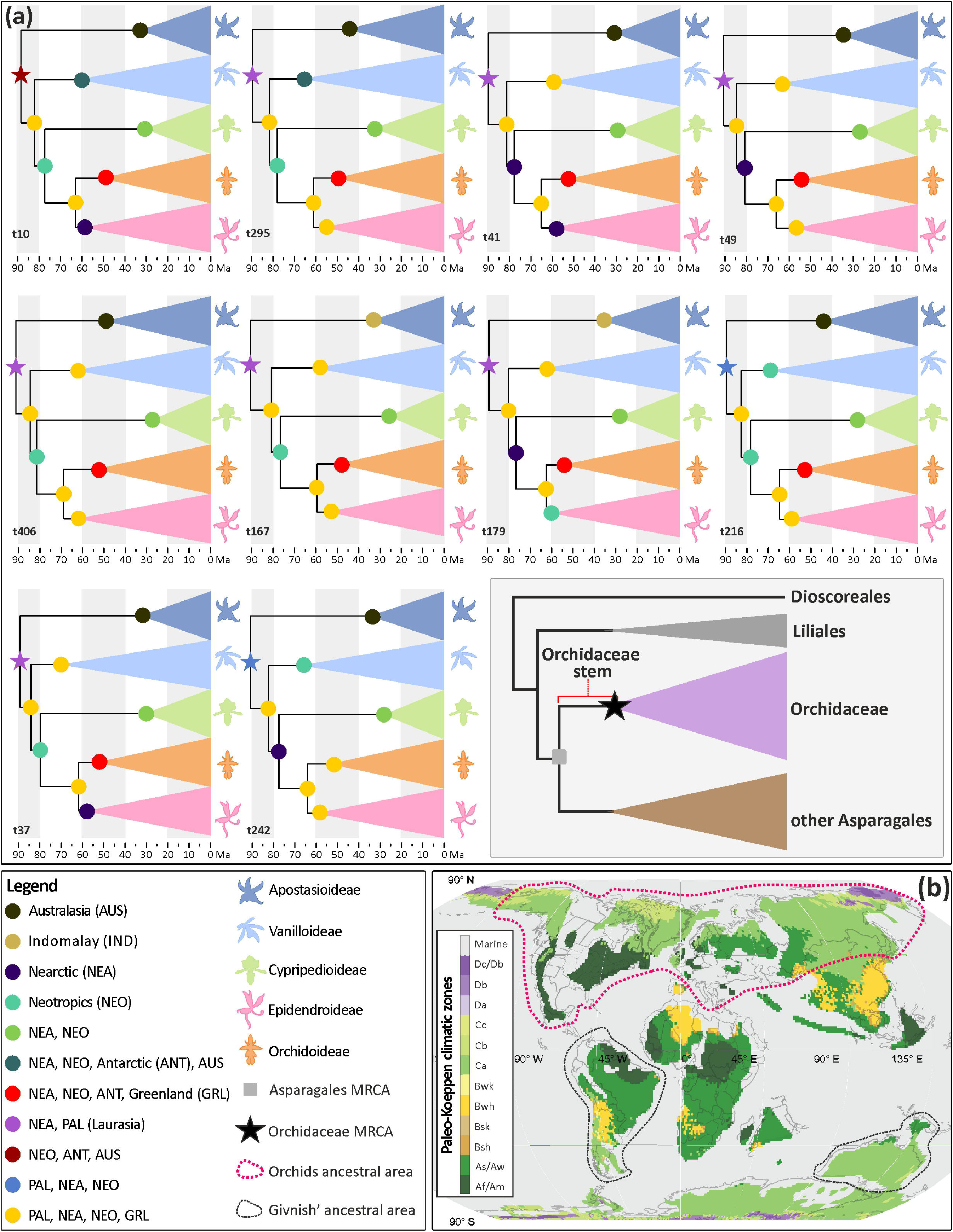
Biogeographic history of the orchid family. A) Ancestral areas at nodes inferred on the ten posterior probability species trees as estimated by a DEC model (see *Methods*) and summarised to the five orchid subfamilies (Inset: A summary of the outgroup sampling considered in our study (the MRCA of orchids is indicated with a black star). B) A palaeoclimatic and tectonic plate reconstruction at 90–80 Ma from **Burgener *et al.*, 2023** showing the possible ancestral range of the orchid MRCA as estimated by **Givnish *et al.*, 2016** and this study. A detailed account of the first and second most likely ancestral areas estimated to the tribe level is provided on Table S5 (complete results including annotated trees with the most likely ancestral area at nodes and alternative ancestral areas, likelihoods and probabilities are available at 10.6084/m9.figshare.22245940).

During the late Cretaceous, Laurasian vegetation was dominated mostly by conifers, and angiosperms, whilst being diverse, were mostly herbs to small trees with early successional strategies restricted to unstable habitats (**Wing **et al.*,* 1998; 2012**). The Laurasian Paleogene has a monsoon-influenced humid subtropical climate that supported closed canopy, broad-leaved deciduous angiosperm forests that extended into high paleolatitudes beyond the Arctic circle (**Eiserhardt **et al.*,* 2017; Korasidis **et al.*,* 2022**).

Proposing an initial diversification of the orchids in Laurasia contrasts with prior inferences of an Australian origin of the family, followed by entry into the Neotropics via Antarctica (**Fig. 4b**, **Givnish **et al.*,* 2016**). **Givnish *et al.*** used nine species as representatives of astelid clades plus one species of Iridaceae, all in Asparagales, most of which are Australian. These outgroups may have influenced the estimate of the Orchidaceae stem node as Australia, for which a ∼40% likelihood was inferred (with Neotropics + Australia inferred as ∼20% likely, and all other ranges together as ∼40%). Their range estimate for the Orchidaceae crown node was Australia ∼20%, Neotropics + Australia ∼30%, and all other ranges added together ∼50%. Our broader sampling of orchid lineages and biogeographic analyses on multiple trees may lead to less phylogenetic and geographic uncertainty (**Rangel **et al.*,* 2015**).

The subfamily Apostasioideae, sister to all other Orchidaceae and comprising only 16 species distributed in Southeast Asia and Northern Australasia (**Li **et al.*,* 2013; Govaerts **et al.*,* 2021**), is inferred by our study to have an ancestral occurrence in Australasia. This would mirror the ranges of numerous other plant groups that were widespread in the Eocene in Eurasia, but which survive only in tropical Southeast Asia today (**Manchester **et al.*,* 2009; Meseguer & Condamine, 2020**). The pattern is known from other 50 groups of gymnosperms and angiosperms (mostly sclerophyllous plants) that formerly occurred in Europe and/or North America – the boreotropical flora of southern Laurasia (**Manchester **et al.*,* 2009**). Descendants from this paleoflora survive today only in eastern Asia, which may have served as a late Cenozoic or Quaternary refugium for these taxa (**Manchester **et al.*,* 2009**). An example amongst Orchidaceae is the subfamily Cypripedioideae (slipper orchids), with 169 extant species (136 Old World and 33 New World). **Guo *et al.*, (2012)** estimated the ancestral range of this clade as Nearctic+Neotropics, in agreement with our study (**Fig. 4**). The Indomalayan region harbours approximately 30% of the extant orchid species, mostly within the Epidendroideae. The oldest known orchid fossil is a hard epidendroid pollinarium found attached to a fungus gnat preserved in Baltic amber, dated to 55-40 Ma (**Poinar & Rasmussen, 2017**), documenting the presence of Epidendroideae at high latitudes in the early Eocene. During this period, evergreen forests covered northern Europe (**Collinson & Hooker, 2003**), and epiphytism likely had already evolved in epidendroid orchids (**Chomicki **et al.*,* 2015**; **Collobert **et al.*,* 2022**). Our biogeographic models, which excluded the Baltic amber pollinarium both as a fossil or as a geographic constraint, inferred that epidendroids were present in the Palearctic around 55–50 Ma, consistent with the fossil evidence. Hard pollinia are prevalent in Epidendroideae and probably appeared early in the history of the subfamily (e.g., in the MRCA of Sobralieae + remainder of epidendroids; **Dressler, 1990; Mosquera-Mosquera **et al.*,* 2019**). However, the evolution of pollinia requires further study, integrating a broader phylogenetic framework in combination with pollinia trait data.

Subsequent colonisation of other biogeographical realms apparently occurred through long-distance dispersals, for example, from Indomalaya to the Afrotropics in tribe Vandeae (Epidendroideae) between 35-10 Ma, or through stepping-stone processes from the Palearctic and Nearctic to the Neotropics in tribe Cymbidieae (Epidendroideae, 30–20 Ma). In contrast, the early diversification of subfamily Orchidoideae probably took place 55-40 Ma between the Neotropics+Nearctic+Antarctic. This was followed by dispersals to Australasia and the Palearctic with subsequent *in-situ* Neotropical diversifications (e.g., Cranichidinae), Nearctic (Spiranthinae), Paleartic, Indomalaya (Goodyerinae) and Afrotropics (e.g., Disinae) (**Data S1**).

### 3.3 A Central American hotspot of orchid speciation

In total, 795,735 records were downloaded from the GBIF and RAINBIO repositories. After rigorously filtering out duplicate records and inaccurate distributions *sensu* the WCVP database (see *Methods*), 495,755 accessions were retained. Analyses of political country species richness indicated that Ecuador, Colombia, and Papua New Guinea are the top three countries in terms of species-richness. Notably, seven out of ten most species-rich countries are located in the Neotropics. Analyses based on the botanical country species richness (as inferred from the WCVP) yielded similar results (**Fig. 5a** [inset], **Fig. S10, Table S6**). An analysis of species richness per grid cell derived from the curated GBIF-RAINBIO dataset, showed that Central America (especially Costa Rica) and the northern Andean region (particularly Ecuador and Colombia) have the highest levels of species richness. These geographical patterns of species richness patterns are thus in agreement with the species richness distributions independently obtained through the WCVP database and support findings of studies conducted at the family level (**Vitt **et al.*,* 2023**) and in the Orchidoideae (**Thompson **et al.*,* 2023**).

**Figure 5.**
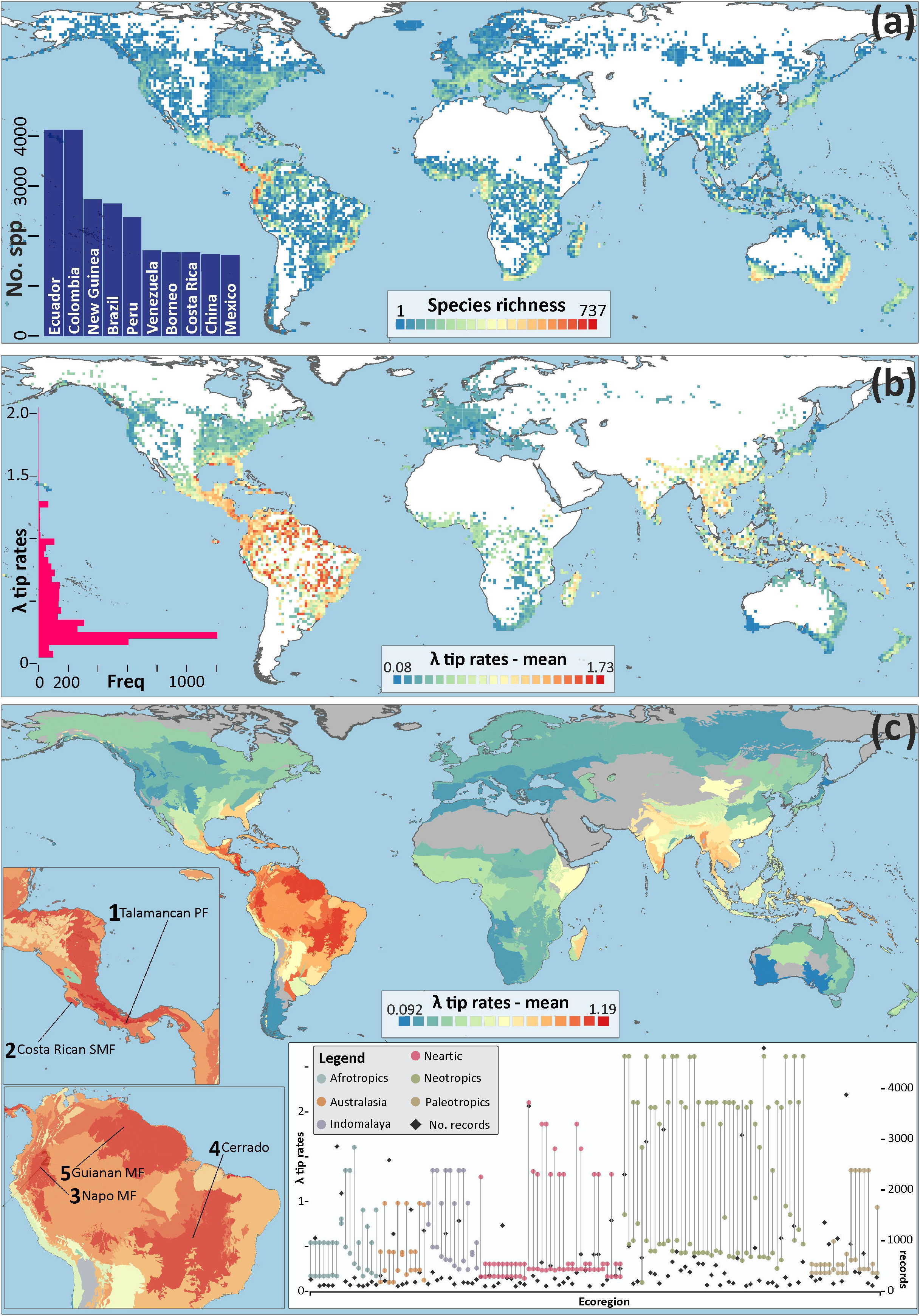
Geography of speciation and species diversity of Orchidaceae. A) Global patterns of species richness per grid-cell (100 ×100 Km), calculated from a curated database of geographical distribution records; reddish colours indicate higher numbers of species per grid-cell whereas bluish colours indicate lower numbers of species per grid-cell. A map of species richness per botanical country is provided in **Figure S10.** (Inset: orchid species numbers reported for the 10 most orchid biodiverse countries). B) Global patterns of mean λ tip rates (linear scale) per grid-cell (100*100 Km) as derived from the BAMM software; warm colours indicate higher λ tip rates grid-cell whereas cold colours indicate lower λ tip rates (Inset: a histogram of the mean λ tip rates attained across all grid cells). C) Global patterns of mean λ tip rates per ecological region, as defined by the WWF, derived from the BAMM software; warm colours indicate higher numbers of mean λ tip rates per ecological region whereas cold colours indicate lower mean λ tip rates. The highlighted geographical areas indicate the five ecoregions with the highest mean λ tip rates (Inset: Maximum and minimum λ tip rate values for ecoregions containing 100 or more geographical distribution records for which λ tip rates were linked).

Geographical speciation (λ) patterns in orchids, as informed from tip λ rates, did not always coincide with the areas of highest current species richness. This discrepancy supports, in some cases, the findings of a recent study on global plant diversification rates and species richness patterns (**Tietje **et al.*,* 2021**). Contrary to the findings of Pérez-Escobar **et al.*,* (**2017a**), the northern Andes do not appear to host the fastest speciating orchid clades in the American tropics despite being one of the most species-rich areas worldwide (**Pérez-Escobar **et al.*,* 2022**). Our findings suggest that southern Mesoamerica, comprising the moist and seasonal forests of Costa Rica and Panama, has the highest speciation rates per grid cell, a region that also exhibits outstandingly high levels of species richness (**Myers **et al.*,* 2000; Mittermeier **et al.*,* 2011; Crain & Fernandez, 2020**, **Fig. 5c; Table S7**). Notably, when considering minimum and maximum values of speciation rates, virtually all ecoregions assessed here presented one- or two-fold variation in their rate (**Fig. 5c, inset**).

Unlike the observed congruence between speciation patterns and species richness throughout most of the Neotropical regions, our study also revealed that southern Australia recovered lower λ tip rates (**Fig. 5c**), even though it shows high levels of species richness (**Myers **et al.*,* 2000**). This implies that elevated species richness in the region might have accumulated at much slower pace and over longer time scales. Whether specific Australian lineages contradict this general pattern remains to be assessed, a task that will require substantially larger sampling. While most of the orchid diversity in southern Australia is terrestrial (**Ackerman, 2019**), a trait associated with lower speciation rates (**Givnish **et al.*,* 2015**), a direct link between high species richness and speciation rates appear to occur in the predominantly terrestrial Orchidoideae (e.g., *Habenaria*; **Thompson **et al.*,* 2023**).

Our speciation rate analyses unveiled multiple accelerations across all subfamilies (**Data S1**). Notably, the majority of these accelerations were nested in the Vanilloideae, Cypripediodeae, Orchidoideae and Epidendroideae subfamilies. Within these, the shifts leading to the fastest tip rates occurred in the Orchidoideae and Epidendroideae, starting from the early Miocene (**Fig. 6a**). These shifts coincided with the initial diversification of species-rich genera such as *Maxillaria and Dendrobium*. In other instances, however, rate increases preceded the divergence of species-rich and depauperate clades as seen with *Lepanthes and Lepanthopsis* or occurred within genera like *Bulbophyllum and Habenaria*. Remarkably, the lineages with the highest speciating lineages are Neotropical epiphytes, including Epidendreae, Maxillariinae, and Oncidiinae. The sole exceptions to this trend are the nearly cosmopolitan terrestrial genus *Habenaria* (**Batista **et al.*,* 2011**) and the genera *Bulbophyllum* and *Dendrobium*, which are mostly epiphytic and distributed mainly in tropical Asia (**Fig. 6b**, **Xiang **et al.*,* 2016; Simpson **et al.*,* 2022**). These findings are in line with previous macroevolutionary studies conducted for the entire family (**Givnish **et al.*,* 2015; Zhang **et al.*,* 2023**) or focused on specific lineages such as Cymbidieae and Pleurothallidinae (**Pérez-Escobar **et al.*,* 2017**) or Orchidinae (**Lai **et al.*,* 2021**).

**Figure 6.**
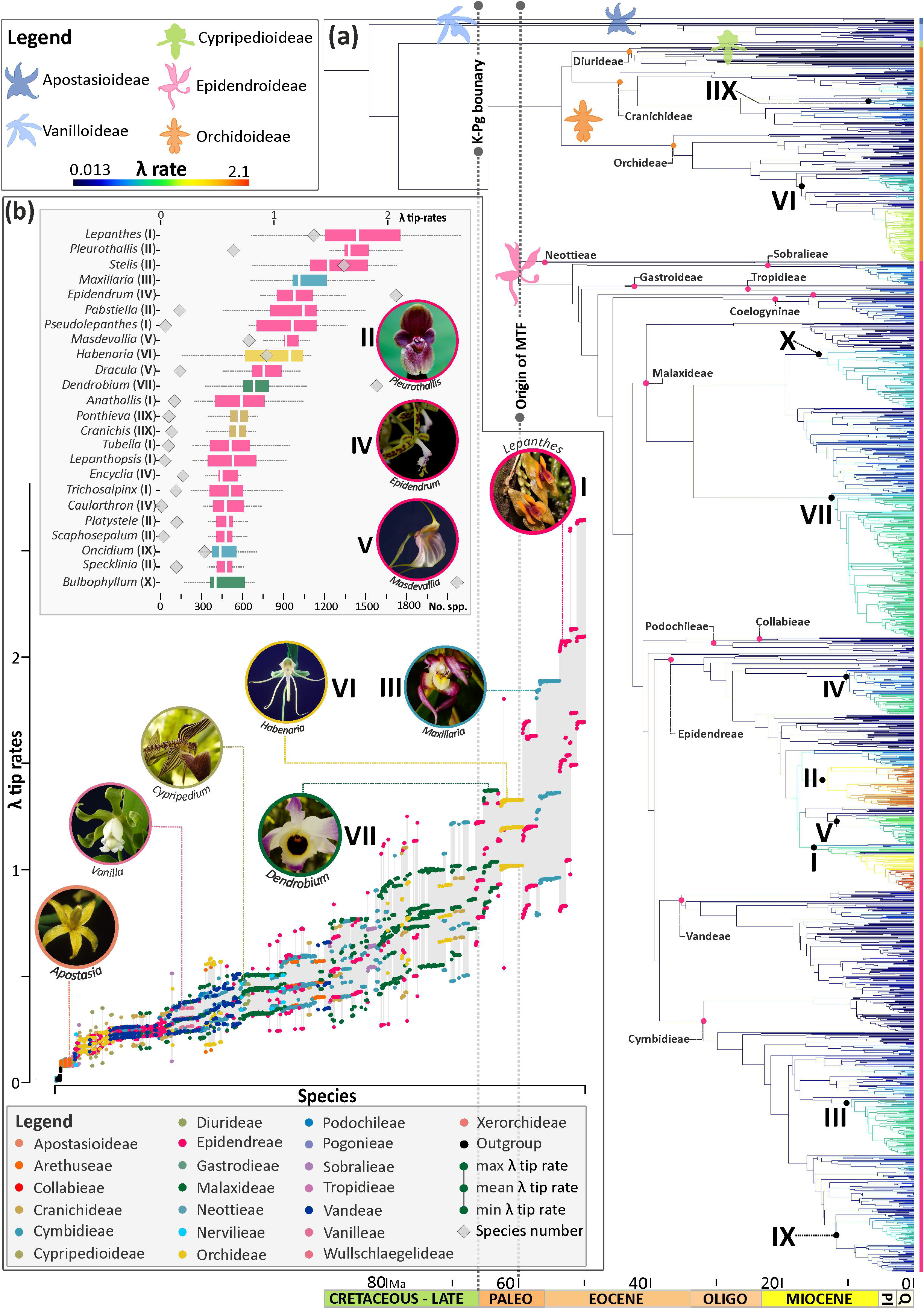
Speciation dynamics of Orchidaceae. A) One of ten posterior probability species trees with modelled speciation (λ) rates along branches (blue colours denote low λ rates, reddish colours indicate high λ rates). Small circles at nodes denote the most recent common ancestors (MRCA) of orchid tribes sampled in this study. Numbers at nodes indicate the lineages with the highest λ tip rates across the entire family. B) Tip λ rate values (maximum, minimum, mean) derived from the ten posterior probability (PP) species trees for all terminals sampled. The dots are colour-coded by taxonomic groups (tribes and subfamilies). (Inset: λ tip rates summarised from the ten PP species trees for the genera associated to the highest λ tip rates [their respective totals of accepted species are provided]. Boxplots are colour-coded by tribes).

Overall, the most rapidly speciating lineages predominantly occur in south-eastern Central America. These lineages span diverse habitats, from lowland dry and wet forests, through cloud forests, to high-elevation grasslands, as exemplified by genera like *Epidendrum*, *Lepanthes*, *Maxillaria*, *Pleurothallis*, and *Stelis* (Fig. **6a**). Interestingly, Central American cloud forests and high-elevation grasslands are of Pliocene origin, making them some of the youngest Neotropical biomes (**Kirby, 2011**). The Late Cenozoic shallow subduction of the young hotspot Cocos ridge under the Panama block (part of the Caribbean plate) is controlling the rapid uplift of the Cordillera de Talamanca in southeast Costa Rica and eastern Panama (**Morell **et al.*,* 2011**). This ongoing process probably started between 5.5 – 3.5 Ma (**Grafe **et al.*,* 2002**), with high elevated mountain uplift rates of ∼1 m per 1 Ma (**Driese **et al.*,* 2007**). The rapid creation of topography within a tropical climate could generate multiple microhabitats that might have enhanced speciation rates. Additionally, the geographic position of Costa Rica and Panama acts as a biological crossroads between the two biodiversity hotspots of northern Mesoamerica and the northern Andes, resulting in the current coexistence of northern and southern lineages (**Burger, 1980; Kapelle, 2016**). These findings support and expand the hypothesis of Central American tropical forests having evolved rapidly and recently (**Cano **et al.*,* 2022**). Nevertheless, although our analyses of geographical speciation rates offer new perspectives on the evolution of orchid diversity, we urge caution in interpreting the relationship between λ and species richness (Tietje *et al.*, 2021). Additionally, our taxon sampling and species-level spatial distributions are likely biased due to uneven collection efforts in certain regions, such as the historically understudied New Guinea (**Camara-Leret **et al.*,* 2020**).

## 4. Conclusions

This study places the initial diversification of orchids in Laurasia during the Late Cretaceous (83 ± 10 Ma) and infers today’s geographic range of Apostasioideae as relictual. The presence of Orchidaceae in Eurasia and North America during the Paleogene, followed by extinction and survival in more southern regions, matches the historical patterns supported by the fossil records of at least 50 genera of other families that were once diverse in Eurasia, but today have smaller surviving groups in tropical Southeast Asia. We reject the hypothesis that Southeast Asia is the region with highest speciation rates (**Givnish **et al.*,* 2016).** Instead, our results show that southern Central America, which contains 4.5% of the world’s flora and fauna in just 0.5% of its land surface, shows the highest speciation rates per grid cell on a global scale. Moreover, these biomes have served as a hotspot for orchid speciation since the Pliocene. Future work could aim at increasing the species-level sampling in phylogenomic trees, particularly for regions and clades currently undersampled. Such additions will further our ability to infer patterns and processes of diversification at increasingly finer spatial and temporal scales.

## 6. Acknowledgments

We thank 3 anonymous reviewers and the editor for valuable feedback which helped us improve this paper. This work was funded by grants from the Calleva Foundation to the Plant and Fungal Trees of Life Project (PAFTOL) at the Royal Botanic Gardens, Kew. OAPE acknowledges support from the Sainsbury Orchid fellowship at the Royal Botanic Gardens Kew and the Swiss Orchid Foundation. The Comisión Institucional de Biodiversidad of University of Costa Rica issued the permit for access to genetic resources under the project B8257. KN acknowledges the Genomics for Australian Plants consortium funded by Bioplatforms Australia (enabled by NCRIS), the Ian Potter Foundation, Royal Botanic Gardens Foundation (Victoria), Royal Botanic Gardens Victoria, the Royal Botanic Gardens and Domain Trust, the Council of Heads of Australasian Herbaria, CSIRO, Centre for Australian National Biodiversity Research and the Department of Biodiversity, Conservation and Attractions, Western Australia. AA acknowledges financial support from the Swedish Research Council (2019-05191), the Swedish Foundation for Strategic Environmental Research MISTRA (Project BioPath), and the Kew Foundation. We thank Sebastian Vieria and Kerry Dressler for granting access to selected photographs of *Apostasia*, *Calochilus*, *Maxillaria* and *Thelymitra*.

## 7. Author contributions

**Conceptualisation:** OAPE, DB, SSR, AA, WJB. **Methodology:** OAPE, FC, DB, SB. **Data curation:** OM, LS, NASP. **Resources**: OAPE, NASP, AZ, DB AQH, BC, KN, MAC, LL. **Software:** OAPE, FC, SB. **Investigation:** NASP, AQH, KA, LS, AZ, LL. **Formal analysis:** OAPE, FC, DB, SB. **Funding acquisition:** WJB, FF, IJL, AA, KN, MAC, OAPE, RPB, RM. **Visualisation:** OAPE, DB, MC, LO, SB. **Writing – original draft:** OAPE, SSR. **Writing – review & editing:** OAPE, SSR, CJ, AA, MC, JA, RT, FLC, DB, NASP, WJB, with further contributions from all co-authors.

## 8. Data availability

The data that support the findings of this study are openly available in https://treeoflife.kew.org/specimen-viewer and at10.6084/m9.figshare.22245940

## 5. References

Ackerman J. 2019. Orchids and the persistent instability principle. Proceedings of the 22th World Orchid Conference.

Ackerman J, Phillips RD, Tremblay RL, Karremans A, Reiter N, Peter CI, Bogarín D, Pérez-Escobar OA, Liu H. 2023. Beyond the various contrivances by which orchids are pollinated: global patterns in orchid pollination biology. Botanical Journal of the Linnean Society in press.

Ali JR, Heaney LR. 2021. Wallace’s line, Wallacea, and associated divides and areas: history of a tortuous tangle of ideas and labels. Biological Reviews 96: 922–942.

Baker WJ, Bailey P, Barber V, Barker A, Bellot S, Bishop D, Botigué LR, Brewer G, Carruthers T, Clarkson JJ, Cook J, Cowan RB, Dodsworth S, Epitawalage N, Francoso E, Gallego B, Johnson M, Kim JT, Leempoel K, Maurin O, Mcginnie C, Pokorny L, Roy S, Malcolm S, Toledo E, Wickett NJ, Zuntini AR, Eiserhardt WL, Kersey PJ, Leitch IJ, Forest F. 2022. Systematic Biology 71(2): 301–319

Balbuena JA, Miguez-Lozano R, Blasco-Costa I. 2013. PACo: a novel Procustres application to cophylogenetic analysis. PLoS ONE 8(4): e61408

Batista J, de Bem Bianchetti L, Gonzalez-Tamayo R, Figueroa XM, Cribb P. 2011. A synopsis of the New World *Habenaria* (Orchidaceae) I. Harvard Papers in Botany 16(1): 1–47.

Beeravolu R, Condamine F. 2016. An extended maximum likelihood inference of geographic range evolution by dispersal, local extinction and cladogenesis. BioRxiv 10.1101/038695

Benzing DH. 2000. Bromeliaceae: profile of an adaptive radiation. Cambridge, UK: Cambridge University Press.

Bouckaert R, Vaughan TG, Barido-Sottani J, S Duchene, M Fourment, Gavryushina A, Heled J, Jones G, Kuhnert D, De Maio N, Matschiner M, Mendes FK, Muller NF, Ogilvie HA, du Plessis L, Popinga A, Rambaut A, Rasmussen D, Siveroni I, Suchard MA, Wu Ch-H, Xie D, Zhang C, Stadler T, Drummond AJ. 2019. BEAST 2.5: an advanced software platform for Bayesian evolutionary analysis. PloS Computational Biology 15(4): e1006650

Bouetard A, Lefeuvre P, Gigant R, Séverine Bory S, Pignal M, Besse P, Grisoni M. 2010. Evidence of transoceanic dispersion of the genus *Vanilla* based on plastid DNA phylogenetic analysis. Molecular Phylogenetics and Evolution 55: 621–630

Brummitt K. 2001. World Geographical Scheme for Recording Plant Distributions. 2^nd^ edition. Pittsburgh, USA: Hunt Institute for Botanical Documentation, Carnegie Mellon University.

Burgener L, Hyland E, Reich RJ, Scotese C. 2023. Cretaceous climates: mapping paleo-Koppen climatic zones using a Bayesian statistical analysis of lithologic, paleontologic and geochemical proxies. Palaeogeography, Palaeoclimatology, Palaeoecology 613: 111373.

Burger, WC. 1980. Why are there so many kinds of flowering plants in Costa Rica? Brenesia 17: 371–88

Camacho C, Coulouris G, Avagyan V, Ma N, Papadopoulos J, Bealer K, Madden TL. 2009. BLAST+: architecture and applications. BMC Bioinformatics 10: 421.

Camara-Leret R, Frodin DG, Adema F, Anderson C, et al. 2020. New Guinea has the world’s richest island flora. Nature 584, 579–583

Cano A, Stauffer FW, Andermann T, Liberal IM, Zizka A, Bacon CD, Lorenzi H, Christe C, Töpel M, Perret M, Antonelli A. 2022. Recent and local diversification of Central American understorey palms. Global Ecology and Biogeography 31: 1513–1525

Chase M. 2001. The origin and biogeography of Orchidaceae. In: Pridgeon AM, Cribb PJ, Chase MW, Rasmussen FN (eds). Genera Orchidacearum: vol. 2. Orchidoideae (part one). Oxford, UK: Oxford University Press.

Chase MW, Cameron KM, Freudenstein JV, Pridgeon AM, Salazar G, van den Berg C, Schuiteman A. 2015. An updated classification of Orchidaceae. Botanical Journal of the Linnean Society 177: 151–174.

Chomicki G, Bidel LPR, Ming F, Coiro M, Zhang X, Wang Y, Jay-Allemand C, Renner SS. 2015. The velamen protects photosynthetic orchid roots against UV-B damage, and a large dated phylogeny implies multiple gains and losses of this function during the Cenozoic. New Phytologist 205(3): 1330–1341.

Christenhusz MJ, Byng JW. 2016. The number of known plant species in the world and its annual increase. Phytotaxa: 261(3): 201–217.

Collinson ME, Hooker JJ. 2003. Paleogene vegetation of Eurasia: framework for mammalian faunas. in: Reumer, J.W.F. & Wessels, W. (eds.): Distribution and migration of tertiary mammals in Eurasia. A volume in honour of Hans de Bruijn. Deinsea 10: 41–83

Collobert G, Perez-Lamarque B, Dubuisson J-Y, Martos F. 2022. Gains and losses of the epiphytic lifestyle in epidendroid orchids: review and new analyses with succulent traits. BioRxiv 10.1101/2022.09.30.510324

Condamine F, Rolland J, Morlon H. 2013. Macroevolutionary perspectives to environmental change. Ecology Letters 16(s1): 72–85.

Conran JG, Bannister JM, Lee DE. 2009. Earliest orchid macrofossils: Early Miocene *Dendrobium* and *Earina* (Orchidaceae: Epidendroideae) from New Zealand. American Journal of Botany 96: 466–474

Couvreur TLP, Forest F, Baker WJ. 2011. Origin and global diversification patterns of tropical rain forests: inferences from a complete genus-level phylogeny of palms. BMC Biology 9:44.

Crain BJ, Fernández M. 2020. Biogeographical analyses to facilitate targeted conservation of orchid diversity in Costa Rica. Diversity and Distributions 26(7): 853–866.

Cribb P, Pridgeon A. 2009. *Claderia*: Phylogenetics. In: Pridgeon AM, Cribb PJ, Chase MW, Rasmussen FN (eds). Genera Orchidacearum: vol. 5. Epidendroideae (part two). Oxford, UK: Oxford University Press.

Dauby G, Zaiss R, Blach-Overgaard A, Catarino L, Damen T, Deblauwe V, Dessin S, Dransfield J, Droissart V, Duarte MC, et al. 2016. RAINBIO: a mega-database of tropical African vascular plants distributions. PhytoKeys 74: 1–18

De Lamotte DF, Fourdan B, Leleu S, Francois L, Clarens P. 2015. Style of rifting and the stages of Pangea break-up. Tectonics 34(5): 1009–1029

Doyle JJ, Doyle JL. 1990. Isolation of plant DNA from fresh tissue. Focus 12: 13–15

Dressler RL. 1990. The orchids: natural history and classification. Cambridge: Harvard University Press.

Driese GS, Kenneth HO, Sally PH, Zheng-Hua L, Debra SJ. 2007. Paleosol evidence for Quaternary uplift and for climate and ecosystem changes in the Cordillera de Talamanca, Costa Rica. Palaeogeography, Palaeoclimatology, Palaeoecology 248: 1–23

Eiserhardt WL, Couvreur TLP, Baker WJ. 2017. Plant phylogeny as a window on the evolution of hyperdiversity in the tropical rainforest biome. New Phytologist 214(4): 1408–1422.

Freudenstein JV, Chase MW. 2015. Phylogenetic relationships in Epidendroideae (Orchidaceae), one of the great flowering plant radiations: progressive specialization and diversification. Annals of Botany 115(4): 665–681

Grace OM, Pérez-Escobar OA, Lucas EJ, Vorontsova MS, Lewis GP, Walker BE, Lohmann LG, Knapp S, Wilkie P, Sarkinen T, Darbyshire I, Lughadha EN, Monro A, Woudstra Y, Demissew S, Muasya AM, Diaz S, Baker WJ, Antonelli A. 2021. Botanical monograph in the Anthropocene. Trends in Plant Science 26(5): 433–441

Kapelle, M. 2016. The montane cloud forests of the Cordillera de Talamanca. In: Kapelle M. Costa Rican Ecosystems. The University of Chicago Press. Chicago.

Givnish TJ, Spalink D, Ames M, Lyon SP, Hunter SJ, Zuluaga A, Iles WJD, Clements MA, Arroyo MTK, Leebens-Mack J, Endara L, Kriebel R, Neubig KM, Whitten WM, Williams NH, Cameron KM. 2015. Orchid phylogenomics and multiple drivers of their extraordinary diversification. Proceedings of the Royal Society B: Biological Sciences 282: 20151553

Givnish TJ, Spalink D, Ames M, Lyon SP, Hunter SJ, Zuluaga A, Doucette A, Giraldo G, McDaniel J, Clements MA, Arroyo MTK, Endara L, Kriebel R, Williams NH, Cameron KM. 2016. Orchid historical biogeography, diversification, Antarctica and the paradox of orchid dispersal. Journal of Biogeography 43(10): 1905–1916.

Górniak M, Paun O, Chase MW. 2010. Phylogenetic relationships within Orchidaceae based on a low-copy nuclear coding gene, *Xdh*: congruence with organellar and nuclear ribosomal DNA results. Molecular Phylogenetics and Evolution 56(2): 784–795.

Govaerts R, Lughadha EN, Black N, Turner R, Paton A. 2021. The World Checklist of Vascular Plants, a continuously updated resource for exploring global plant diversity. Scientific Data 8: 215

Grafe KW, Frisch IM, Villa MM. 2002. Geodynamic evolution of southern Costa Rica related to low-angle subduction of the Cocos Ridge: constraints from thermochronology. Tectonophysics 348: 187– 204

Gustafsson ALS, Verola CF, Antonelli A. 2010. Reassessing the temporal evolution of orchids with new fossils and a Bayesian relaxed clock, with implications for the diversification of the rare South American genus *Hoffmannseggella* (Orchidaceae: Epidendroideae). BMC Evolutionary Biology 10(1): 1–13

Guo Y-Y, Luo Y-B, Liu Z-J, Wang X-Q. 2012. Evolution and biogeography of the slipper orchids: eocene vicariance of the conduplicate genera in the Old and New World Tropics. PLoS ONE 7(6): e38788. doi:10.1371/journal.pone.0038788

Huson DH, Bryant D. 2006. Application of phylogenetic networks in evolutionary studies. Molecular Biology and Evolution 23(2): 254–267.

Johnson MG, Gardner EM, Liu Y, Medina R, Goffinet B, Shaw AJ, Zerega NJ, Wicket NJ. 2016. HybPiper: extracting coding sequence and introns for phylogenetics from high-throughput sequencing reads using target enrichment. Applications in Plant Sciences 4(7): 1600016

Johnson MG, Pokorny LP, Dodsworth SD, Botigué LR, Cowan RS, Devault A, Eiserhardt WL, Epitawalage N, Forest F, Kim JT, Leebens-Mack JH, Leitch IJ, Maurin O, Soltis DE, Soltis PS, Wong G, Baker WJ, Wickett NJ. 2019. A universal probe set for targeted sequencing of 353 nuclear genes from any flowering plant designed using k-medoids clustering. Systematic Biology 68(4): 594–606.

Jones DL. 1997. *Cooktownia robertsii*, a remarkable new genus and species of Orchidaceae from Australia. Austrobaileya 5(1): 71–78.

Katoh K, Standley DM. 2013. MAFFT: multiple sequence alignment software version 7: improvements in performance and usability. Molecular Biology and Evolution 30: 772–780.

Kirby SH. 2011. Active mountain building and the distribution of “core” Maxillariinae species in tropical Mexico and Central America. Lankesteriana 11(3): 275–291.

Korasidis VA, Wing SL, Shields CA, Kiehl JT. 2022. Global changes in terrestrial vegetation and continental climate during the Paleocene-Eocene Thermal Maximum. Paleoceanography and Paleoclimatology 37: e2021PA004325.

Lai Y-J, Han Y, Schuiteman A, Chase MW, Xu S-Z, Li J-W, Wu J-Y, Yang B-Y, Jin X-H. 2021. Diversification in Qinghai-Tibet Plateau: Orchidinae (Orchidaceae) clades exhibiting pre-adaptations play a critical role. Molecular Phylogenetics and Evolution 157: 107062

Li Y, Ma L, Liu D-K, Zhao X-W, Z D, Ke S, Chen G-Z, Zheng Q, Liu Z-J, Lan S. 2023. *Apostasia fujianica* (Apostasioideae, Orchidaceae), a new Chinese species: evidence from morphological, genome size and molecular analyses. Phytotaxa 583(3): 277–284.

Louca S, Pennell MW. 2021. Why extinction estimates from extant phylogenies are so often zero. Current Biology 31(14): 3168–3173.

Magallón S, Sánchez-Reyes LL, Gómez-Acevedo SL. 2018. Thirty clues to the exceptional diversification of flowering plants. Annals of Botany 123: 491–503.

Maldonado C, Molina CI, Zizka A, Persson C, Taylor CM, Alban J, Chilquillo E, Ronsted N, Antonelli A. 2015. Estimating species diversity and distribution in the era of Big Data: to what extent can we trust public databases? Global Ecology and Biogeography 24(8): 973–984

Manchester SR, Chen Z-D, Lu A-M, Uemura K. 2009. Eastern Asian endemic seed plant genera and their paleogeographic history throughout the northern hemisphere. Journal of Systematics and Evolution 47: 1–42.

Matzke NJ. 2013. Probabilistic historical biogeography: new models for founder-event speciation, imperfect detection, and fossil allow improved accuracy and model-testing. Frontiers of Biogeography 5:243–248.

Meseguer AS, Condamine FL. 2020. Ancient tropical extinctions at high latitudes contributed to the latitudinal diversity gradient. Evolution 74(9): 1966–1987.

Mittermeier RA, Turner WR, Larsen FW, Boorks TM, Gascon C. 2011. Global biodiversity conservation: the critical role of hotspots. In: Zachos FE, Habel JC. Biodiversity hotspots: distribution and protection of conservation priority areas. Heidelberg, Germany: Springer.

Morell KD, Fisher DM, Gardner TW, La Femina P, Davidson D, Teletzke A. 2011. Quaternary outer fore-arc deformation and uplift inboard of the Panama Triple Junction, Burica Peninsula. Journal of Geophysical Research 116: B05402.

Mosquera-Mosquera HR, Valencia-Barrera RM, Acedo C. 2019. Variation and evolutionary transformation of some characters of the pollinarium and pistil in Epidendroideae (Orchidaceae). Plant Systematics and Evolution 305: 353–374.

Myers N, Mittermeier RA, Mittermeier CG, da Fonseca GAB, Kent J. 2000. Biodiversity hotspots for conservation priorities. Nature 403(24): 853.

Nauheimer L, Schley RJ, Clements MA, Micheneau C, Nargar K. 2018. Australian orchid biogeography at continental scale: molecular phylogenetic insights from the sun orchids (*Thelymitra*, Orchidaceae). Molecular Phylogenetics and Evolution 127: 304–319.

Niissalo MA, Leong PKF, Tay FEL, Choo LM, Kurzweil H, Khew GS. 2023. A new species of *Claderia* (Orchidaceae). Gardens’ Bulletin Singapore 75(1): 21–41.

Pérez-Escobar OA, Balbuena JA, Gottschling M. 2016. Rumbling orchids: how to assess divergent evolution between chloroplast endosymbionts and the nuclear host. Systematic Biology 65(1): 51–65.

Pérez-Escobar OA, Chomicki G, Condamine FL, Karremans AP, Bogarín D, Matzke NJ, Silvestro D, Antonelli A. 2017a. Recent origin and rapid speciation of Neotropical orchids in the world’s richest plant biodiversity hotspot. New Phytologist 215(2): 891–905.

Pérez-Escobar OA, Gottschling M, Chomicki G, Condamine FL, Klitgaard B, Pansarin E, Gerlach G. 2017b. Andean mountain building did not preclude dispersal of lowland epiphytic orchids in the Neotropics. Scientific Reports 7(1): 4919.

Pérez-Escobar OA, Dodsworth S, Bogarín D, Balbuena JA, Schley RJ, Kikuchi IZ, Morris SK, Epitawalage N, Cowan R, Maurin O, Zuntini A, Arias T, Serna-Sánchez A, Gravendeel B, Torres MF, Nargar K, Chomicki G, Chase MW, Leitch IJ, Forest F, Baker WJ. 2021a. Hundreds of nuclear and plastid loci yield novel insights into orchid relationships. American Journal of Botany 108(7): 1166–1180.

Pérez-Escobar OA, Bellot S, Przelomska NAS, Flowers JM, Nesbitt M, Ryan P, et al. 2021b. Molecular clocks and archaeogenomics of a late period Egyptian date palm leaf reveal introgression from wild relatives and add timestamps on the domestication. Molecular Biology and Evolution 38(10): 4475–4492.

Pérez-Escobar OA, Zizka A, Bermúdez MA, Meseguer AS, Condamine FL, Hoorn C, Hooghiemstra H, Pu Y, Bogarín D, Boschman LM, Pennington RT, Antonelli A, Chomicki G. 2022. The Andes through time: evolution and distribution of Andean floras. Trends in Plant Science 27(4): 1–12.

Poinar G Jr. 2016a. Orchid pollinaria (Orchidaceae) attached to stingless bees (Hymenoptera: Apidae) in Dominican amber. Neues Jahrbuch fuDr Geologie und Paläontologie – Abhandlungen 279: 287–293.

Poinar G Jr. 2016b. Beetles with orchid pollinaria in Dominican and Mexican amber. American Entomologist 62: 172–177.

Poinar G, Rasmussen FN. 2017. Orchids from the past, with a new species in Baltic amber. Botanical Journal of the Linnean Society 183: 327–333.

Portik DM, Wiens JJ. 2020. SuperCRUNCH: a bioinformatics toolkit for creating and manipulating supermatrices and other large phylogenetic datasets. Methods in Ecology and Evolution 11(6): 7763–772.

Rabosky DL, Santini F, Eastman J, Smith SA, Sidlauskas B, Chang J, Alfaro ME. 2013. Rates of speciation and morphological evolution are correlated across the largest vertebrate radiation. Nature Communications 4(1):1958.

Rabosky DL, Grundler M, Anderson C, Title P, Shi JF, Brown JW, Huang H, Larson JG. 2014. BAMM tools: an R package for the analysis of evolutionary dynamics on phylogenetic trees. Methods in Ecology and Evolution 5(7): 701–707.

Ramírez SR, Gravendeel B, Singer RB, Marshall CR, Pierce NE. 2007. Dating the origin of the Orchidaceae from a fossil orchid with its pollinator. Nature 448: 1042–1045.

Rangel TF, Colwell RK, Graves GR, Fucikova K, Rahbek C, Diniz-Filho JF. 2015. Phylogenetic uncertainty revisited: implications for ecological analyses. Evolution 69(5): 1301–1312.

Ree RH, Smith SA. 2008. Maximum likelihood inference of geographic range evolution by dispersal, local extinction, and cladogenesis. Systematic Biology 57(1): 4–14.

Revell LJ. 2012. phytools: an R package for phylogenetic comparative biology (and other things). Methods in Ecology and Evolution 3: 217– 223.

Serna-Sánchez M, Pérez-Escobar OA, Bogarín D, Torres-Jimenez MF, Alvarez-Yela AC, Arcila-Galvis JE, Hall C, de Barros D, Pinheiro F, Dodsworth S, Chase MW, Antonelli A, Arias T. 2021. Plastid phylogenomics resolves ambiguous relationships within the orchid family and provides a solid timeframe for biogeography and macroevolution. Scientific Reports 11: 6858

Simpson L, Clements MA, Orel HK, Crayn DM, Nargar K. 2022. Plastid phylogenomics clarifies broad-level relationships in *Bulbophyllum* (Orchidaceae) and provides insights into range evolution of Australasian section *Adelopetalum*. BioRxiv 10.1101/2022.07.24.500920

Smith SA, Moore MJ, Brown JW, Yang Ya. 2015. Analysis of phylogenomic datasets reveal conflict, concordance, and gene duplications with examples from animals and plants. BMC Evolutionary Biology 15: 150

Smith SA, Brown JW, Walker JF. 2018. So many genes, so little time: a practical approach to divergence-time estimation in the genomic era. PLoS ONE 13(5): e0197433.

Soreng RJ, Peterson PM, Romaschenko K, Davidse G, Zuloaga FO, Judziewicz EJ, Filgueiras TS, Davis JI, Morrone O. 2015. A worldwide phylogenetic classification of the Poaceae (Gramineae). Journal of Systematics and Evolution 53(2): 117–137

Stamatakis, A. 2014. RAxML version 8: A tool for phylogenetic analysis and post-analysis of large phylogenies. Bioinformatics 30: 1312–1313.

Stull GW, Pham KK, Soltis PS, Soltis DE. 2023. Deep reticulation: the long legacy in vascular plant evolution. The Plant Journal doi10.1111/tpj.16142.

Thompson JB, Davis KE, Dodd HO, Priest NK. 2023. Speciation across the Earth driven by global cooling in terrestrial orchids. Proceedings of the National Academy of Sciences 120: e2102408120

Tietje M, Antonelli A, Baker WJ, Govaerts R, Smith SA, Eiserhardt WL. 2022. Global variation in diversification rate and species richness are unlinked in plants. *Proceedings of the National Academy of Sciences*, USA 119(27): e2120662119

Title PO, Rabosky DL. 2019. Tip rates, phylogenies and diversification: what are we estimating, and how good are the estimates? Methods in Ecology and Evolution 10(6): 821–834.

Töpel M, Zizka A, Maria Fernanda Calió MF, Scharn R, Silvestro D, Antonelli A. 2017. SpeciesGeoCoder: fast categorization of species occurrences for analyses of biodiversity, biogeography, ecology, and evolution. Systematic Biology 66(2):145–151.

Walker B. 2022. kewr: R Package to Access Kew Data APIs. ttps://barnabywalker.github.io/kewr/, https://github.com/barnabywalker/kewr/

Weins JJ, Donoghue MJ. 2004. Historical biogeography, ecology and species richness. Trends in Ecology and Evolution 19: 639–644.

Wing SL, Boucher LD. 1998. Ecological aspects of the Cretaceous flowering plant radiation. Annual Reviews of Earth and Planetary Science 26: 379–421.

Wing SL, Strömberg C, Hickey LJ, Tiver F, Willis B, Burnham RJ, Behrensmeyer AK. 2012. Floral and environmental gradients on a Late Cretaceous landscape. Ecological Monographs 82: 23–457

Xiang X-G, Mi X-Ch, Zhou H-L, Li J-W, Chung Sh-W, Li D-Z, Huang W-Ch, Jin W-T, Li Z-Y, Huang L-Q, Jin X-H. 2016. Biogeographical diversification of mainland Asian *Dendrobium* (Orchidaceae) and its implications for the historical dynamics of evergreen broad-leaved forest. Journal of Biogeography 43(7): 1310–1323.

Van den Berg C, Goldman DH, Freudenstein JV, Pridgeon AM, Cameron KM, Chase MW. 2005. AN overview of the phylogenetic relationships within Epidendroideae inferred from multiple DNA regions and recircumscription of Epidendreae and Arethuseae (Orchidaceae). American Journal of Botany 92(4): 613–624.

Van der Niet T, Linder PH. 2008. Dealing with incongruence in the quest of the species tree: a case of study from the orchid genus *Satyrium*. Molecular Phylogenetics and Evolution 47: 154–174.

Velasco JA, Pinto-Ledezma JN. 2022. Mapping species diversification metrics in macroecology: prospects and challenges. Frontiers in Ecology and Evolution 10:1–18.

Viruel J, Segarra-Moragues JG, Raz L, Forest F, Wilkin P, Sanmartín I, Catalán P. 2015. Late Cretaceous - Early Eocene origin of yams (*Dioscorea*, Dioscoreaceae) in the Laurasian Palaeartic and their subsequent Oligocene-Miocene diversification. Journal of Biogeography 43(4): 750–672.

Vitt P, Taylor A, Rakosy D, Kreft H, Meyer A, Wigelt P, Knight TM. 2023. Global conservation prioritization of the Orchidaceae. Scientific Reports 13: 6718

Zhang C, Rabiee M, Sayyari E, Mirarab S. 2018. ASTRAL-III: polynomial time species tree reconstruction from partially resolved gene trees. BMC Bioinformatics 19:153.

Zhang C, Zhao Y, Braun EL, & Mirarab S. (2021). TAPER: Pinpointing errors in multiple sequence alignments despite varying rates of evolution. Methods in Ecology and Evolution, 00, 1– 14.

Zhang G, Hu Y, Huang M-Z, Huang W-C, Liu D-K, Zhang D, Hu H, Downing JL, Liu Z-J, Ma H. 2023. Comprehensive phylogenetic analyses of Orchidaceae using nuclear genes and evolutionary insights into epiphytism. Journal of Integrative Plant Biology, doi10.1111/jipb.13462

