## Supplemental Material for "The Origin And Speciation Of Orchids"

^1^Royal Botanic Gardens, Kew, TW9 3AE, London, UK.

^2^Lankester Botanical Garden, University of Costa Rica, Cartago, Costa Rica.

^3^Naturalis Biodiversity Centre, Leiden, The Netherlands.

^4^University of Portsmouth, PO1 2DY, Portsmouth, UK.

^5^University of Puerto Rico – Rio Piedras, Puerto Rico, USA.

^6^ICBiBE, Universitat de València, Valencia, Spain.

^7^Swiss Orchid Foundation, Basel, Switzerland.

^8^Jardín Botánico Rafael Maria Moscoso, Santo Domingo, Dominican Republic.

^9^Jose Celestino Mutis Botanic Garden, Bogota, Colombia.

^10^University of Ioannina, GR 45110, Ioannina, Greece.

^11^Durham University, DH13LE, Durham, UK.

^12^Centre for Australian National Biodiversity Research (joint venture between Parks Australia and CSIRO), GPO Box 1700, Canberra, ACT 2601, Australia

^13^Universidad Pontificia Javeriana, Seccional Cali, Colombia.

^14^Herbarium AMO, Mexico City, Mexico.

^15^Singapore Botanic Gardens, 1 Cluny Road, Singapore.

^16^Smithsonian Tropical Research Institute, Apartado 0843-03092, Panama City, Panama.

^17^Universidade Federal do Paraná, Brazil

^18^Reserva Biológica Guaitil, Costa Rica.

^19^National Research Collections Australia, Commonwealth Industrial and Scientific Research Organisation (CSIRO), GPO Box 1700, Canberra, ACT 2601, Australia

^20^Universidad Nacional Autónoma de México, Mexico City, Mexico.

^21^Australian Tropical Herbarium, James Cook University, GPO Box 6811, Cairns, QLD 4878

^22^Instituto Politécnico Nacional, CIIDIR unidad Oaxaca, Mexico.

^23^University of Sheffield, Sheffield, UK.

^24^Universidade Estadual de Feira de Santana, Brazil

^25^Universidad del Valle, Valle del Cauca, Colombia.

^26^Department of Environment and Agriculture, Curtin University, Perth, WA 6102, Australia

^27^Institut des Sciences de l’Evolution de Montpellier (Université de Montpellier | CNRS | IRD | EPHE), Place Eugène Bataillon, Montpellier, France.

^28^Scientific Research Organisation (CSIRO), GPO Box 1700, Canberra, ACT 2601, Australia

^29^Washington University, St Louis, USA.

^30^Gothenburg Global Biodiversity Centre, Department of Biological and Environmental Sciences

^31^University of Gothenburg, Gothenburg, Sweden

^32^Wuhan Botanical Garden, Chinese Academy of Sciences, Wuhan, China

^33^Department of Biology, University of Oxford, Oxford, UK

^*^Lead authors

^§^Senior authors

The following Supporting Information is available for this article:

**Supplementary Methods S1**

**Supplementary Results and Discussion S1**

**Supplementary Figures S1-S11**

**Supplementary Tables S1-S7**

**Supplementary Data S1**

**Supplementary Methods S1**

***DNA library preparation, sequencing, and data analysis***

DNA extraction was conducted from 1.5-4 mg of silica-dried fresh tissue or herbarium specimen tissue, constituting leaf or occasionally flower. We used a sterile steel bead for mechanical disruption in a SPEX® sample prep tissue homogeniser (SPEX Inc, NJ, USA), applying two to four disruption cycles of 1 min at 1350Hz. Next, we added 750µL CTAB with 2% v/v 2-mercaptoethanol (Doyle & Doyle, 1990) for a 30-min incubation at 65°C, followed by either a 4-hour incubation at 57°C (silica-dried samples) or an overnight incubation at 57°C (herbarium specimens). We then added 750 µL SEVAG (24:1 chloroform:isoamyl alcohol), placed the samples on a shaking incubator for 30 mins, and centrifuged the samples to induce phase separation. Next, we separated off the upper phase containing the DNA fraction and added to this a 0.7× volume of isopropanol for a 4–5-day incubation at -20°C to encourage DNA precipitation. Finally, we subjected the samples to two rounds of centrifugation and supernatant removal, firstly to remove the isopropanol and secondly to wash the DNA pellet with 75% ethanol. Subsequently, the DNA pellet was eluted into 70µL 10mM Tris-Cl and purified using a paramagnetic bead clean-up method with a 2:1 ratio of Ampure XP beads (Beckman Coulter, Brea, CA, USA) to DNA elution. To gauge the size distribution of the genomic DNA and, on this basis, determine which samples might need shearing before library preparation, we visualised the samples on a 1% agarose gel.

Using 18-200 ng starting material, we prepared DNA libraries using NEB Next Ultra II Library Prep Kits (New England Biolabs, Ipswich, MA, USA), according to the manufacturer’s protocol, but with half-volume reactions. Where possible (silica dried material and herbarium specimens with DNA of higher integrity), we aimed for insert sizes of ~350bp, by shearing the DNA with a Covaris ME220 Focussed Ultrasonicator (Covaris LLC, Woburn, MA, USA). The libraries were indexed with NEBNext Multiplex Oligos for Illumina (New England Biolabs, Ipswich, MA, USA) and amplified using 6-14 PCR cycles. The yield was estimated using a Quantus fluorometer (Promega, Madison, WI, USA), and fragment size distribution was estimated using either a 4200 TapeStation system or an Agilent 2100 BioAnalyser (Agilent Technologies, Santa Clara, CA, USA). Finally, we used the Angiosperms353 probe set to enrich these genomic libraries for 353 low-copy nuclear genes (Johnson et al., 2019), modifying the equimolar pooling of libraries into groups of 10-20 libraries per capture reaction and total input of up to 1µg. Furthermore, for the herbarium samples, we applied a lowered hybridisation temperature – of 62°C and a prolonged hybridization time – of 40 hours. The sequencing of paired-end genomic libraries (150 bp × 2) was conducted on an Illumina HiSeq by Macrogen (Geumcheon, South Korea). Newly generated Illumina sequencing data and associated voucher information are freely available via the Kew Tree of Life Explorer (https://treeoflife.kew.org/).

We retrieved the Angiosperms353 coding loci from the quality trimmed and filtered reads and a refined target file produced for the orchid family (McLay et al., 2021) using the pipeline HybPiper v.2.1.6 (Johnson et al., 2016) and the following parameters: *a*) coverage cut-off of 8x for *de-novo* assembly; b) the BWA program v.0.7 for read mapping (Li and Dublin 2009). We excluded all paralog genes flagged by the pipeline (5 genes).

***Gene tree incongruence analysis***

Gene tree incongruence is a pervasive phenomenon in plant lineages (e.g., Vargas et al. 2017; McLay et al. 2023), including orchids (e.g., Van der Niet and Linder 2008; Pérez-Escobar et al. 2017). To assess whether our nuclear and plastid Sanger phylogenies are incongruent, we conducted a goodness-of-fit test (PACo: Balbuena et al., 2013) that has been widely implemented in multiple plant lineages (e.g., Perez-Escobar et al., 2016, Vargas et al., 2017). This test quantifies the level of discordance between any two given tree topologies and explicitly tests that the similarities between these topologies are not higher than expected by chance by comparing their Euclidean distances obtained from distance. Using ML bipartition trees independently produced from the ITS and *mat*K alignments (see *Methods* section), we conducted this goodness-of-fitness test by implementing 100 permutations in the software R (R Core Team 2022). Additionally, we computed the proportion of gene tree quartets that satisfied the species tree topology by conducting a multispecies coalescent (MSC) analysis using the software ASTRAL-III (Zhang et al., 2018) and ML bipartition trees with branches having a likelihood bootstrap support < 20% collapsed.

***Molecular clock dating analyses and species-level phylogeny assembly***

For the absolute age estimation analysis of the top 25 low-copy nuclear genes, we included a single representative per genus (i.e., the sample for which genes the most were retrieved) except for 13 genera (*Aeranthes*, *Agrostophyllum*, *Brachionidium*, *Cleisocentron*, *Cleisostoma*, *Clowesia, Diodonopsis, Dipodium, Octomeria, Restrepia, Trachoma, Trichocentrum, Trichoglottis*, *Tubella*) for which we could not obtain Sanger sequences that passed our filtering procedure. For these groups, we retained the original species sampling.

We constructed multiple ultrametric species trees instead of relying on a single consensus maximum credibility tree to best account for branch length variation and uncertainty around poorly supported phylogenetic relationships during the ancestral area and speciation rate estimation. We first randomly sampled 10 posterior trees derived from the BEAST analyses conducted on the genus-level Angiosperms353 and the ITS-*mat*K Sanger species-level datasets. Then, for every genus represented in the Angiosperms353 chronograms, we pruned its counterpart clade sampled on the species-level Sanger chronogram and proceeded to graft it onto the corresponding stem of the Angiosperms353 chronogram. Only monophyletic groups from the Sanger chronogram were grafted into the Angsioperms353 tree. Additionally, we also checked for potential incongruences between the stem age and the crown node of the Angiosperm353 and Sanger chronograms, respectively (see *Supplementary Results and Discussion S1*). Here, whenever we encountered discordance between these ages, we proceeded to rescale the age of the genus’ MRCA in the Sanger tree prior to grafting, by using the NGS genus’ 95% High Probability Density (HPD) lower bound value (see Figure 2 in main text). To achieve this, we relied on the function *chronos* of the R package APE (Paradis et al., 2004). This operation was conducted on each pair of randomly sampled posterior trees (hence called PP species trees). To assess the robustness of the phylogenetic relationships derived from the PP species trees, we repeated the pruning and grafting procedure in 500 randomly sampled genus-level Angiosperm353 and species-level Sanger MCMC trees and we produced a maximum clade credibility tree (MCC tree) from the post-burn-in (10%) posterior probability trees using TreeAnnotator v.2.6 (available at <https://www.beast2.org/treeannotator/>). We then counted the proportion of bipartitions falling on different support intervals (**Figure S8**).

**Supplementary Results and Discussion S1**

***Phylogenetic relationships in the Orchidaceae***

Building a comprehensively sampled and robust orchid tree of life is a challenging task. Beyond the quantitative challenge of comprising hundreds of recognized genera, there is a myriad of oft-endemic species with narrow distribution ranges, which renders these species challenging to account for in taxon samplings. Our Multispecies coalescent tree (MSC) and Maximum likelihood (ML) analyses produced the most densely sampled, well-supported, to-date genus-level phylogenomic framework for the Orchidaceae (**Figure 2**). Here, *in silico,* gene capture efficiency was similar to that reported by Pérez-Escobar et al. (2021), ranging from 4 to 90% per sequenced sample (**Supp. Fig. S1-S6**). Individual gene and concatenated ML analyses conducted on the Angisoperms353 low copy nuclear gene alignments indicated that bipartitions frequently attained moderate (70%) to maximum (90-100%) likelihood bootstrap support values (LBS) (**Figure 2A,B**). In addition, the MSC tree analysis revealed that topological incongruence between the Angisoperms353 low-copy nuclear gene trees was minimal (normalized Q-score: 0.96; Figure 2C). This suggests that factors known to drive tree discordance, such as hybridisation, incomplete lineage sorting, and paralogy (**Pérez-Escobar et al., 2016**) did not substantially influence relationships in the Orchidaceae as inferred from our datasets. Interestingly, a previous MSC analysis conducted on a reduced set of ML gene trees (249) and genera (89) yielded a lower normalized Q-score (0.89; **Pérez-Escobar et al., 2021**), indicating that taxon sampling could affect gene tree congruence (**van der Niet & Linder 2008**).

The test of topological congruence between the ITS and *mat*K bipartition trees conducted in PACo revealed that both topologies are largely congruent (***m^2^_xy_*** = 35.76235, P <0.0001). In addition, we also investigated the proportion of ITS and *mat*K gene tree quartets that match the species tree in ASTRAL-III. The analysis indicated that 99.1% of the gene tree quartets agree with the species tree topology (normalised quartet score = 0.9917), in line with the results obtained by PACo. Both analyses indicate that incongruence between the ITS and *mat*K DNA matrices is limited, and therefore, they are suitable for concatenation and phylogenetic inference.

An inspection of the local quartet values derived from the MSC analyses indicated that between 40 and 100% of the gene tree quartets agreed with the species tree relationships, without an obvious link to phylogenetic depth. The bipartitions linked to quartet values < 40 were associated with samples whose gene recovery was poor (e.g., *Cyrtostylis huegelii* Endl.: ~10% recovery; *Bifrenaria aurantiaca* Lindl.: ~25%; *Aphyllorchis queenslandica* Dockrill: ~25%), or to relationships that have historically been difficult to resolve (e.g., MRCA of Neottieae+remainder of Epidendroids, MRCA of Cymbideae+Epidendreae; **Chase et al., 2015**). In any case, the proportion of gene tree quartets that agreed with a particular bipartition recovered by the species tree was often higher than any other second most frequent alternative bipartition (Supp. Fig. S1-S8). The few exceptions to this pattern were: *a*) the branch leading to the MRCA of Caladeniinae, Acianthinae, and Prasophyllinae in the Orchidoideae, three clades whose relationships were not interrogated through high-throughput sequencing datasets and remained unresolved (**Kores et al., 2001; Chase et al., 2015**); *b*) bipartitions nested inside *Cycnoches* Lindl. (Catasetinae, Cymbidieae) and *Pleurothallis* R.Br. (Pleurothallidinae, Epidendreae)*,* two genera derived from rapid diversifications that are known to have experienced interspecific hybridisation leading to gene-tree conflicts (**Pérez-Escobar 2016; Mauad et al., 2021**).

Relationships derived from our extensive taxon sampling (Figure 2A) largely agreed with previously published studies (**Li et al., 2019;** **Serna-Sanchez et al., 2021**), yet only some groups thought to be monophyletic were not recovered as natural groups. These include the orchidoid subtribes Drakaeinae+Megastylidinae, the epidendroid tribe Neottieae, and subtribes Adrorhizinae+Polystachyinae and Calypsoinae. In all cases, the new relationships fell within moderately to strongly supported bipartitions in both concatenated ML and MSC analyses (LBS > 75 and local quartet scores >=35). More recently, **Zhang et al., (2023)** produced a broader phylogenomic framework of the Orchidaceae, using a range of 610 to 1,195 nuclear ortholog genes. Our phylogenies are in close agreement with those of **Zhang et al., (2023)** yet a small number of discordances, which appear to be important, were found in subtribal and generic relationships and that have never been recovered in previous studies (e.g., Gorniak et al., 2010; Serna-Sánchez et al., 2021). Apparently, no approach to quantify the proportion of gene tree conflict in Zhang’s et al. (2023) dataset was implemented, rendering it difficult to assess the credibility of the proposed novel relationships. Assessing gene tree incongruence during phylogenetic inference is crucial for accurate species tree estimation. This is particularly true for angiosperms, since topological discordance is a pervasive phenomenon across many lineages (Smith et al., 2015; Stull et al., 2023), including orchids (**van der Niet and Linder, 2008; Pérez-Escobar et al. 2016; 2017**).

The novel relationships revealed by the inclusion of more generic clades and previously unsampled genera suggest that more research needs to be done to obtain a comprehensive understanding of the phylogenetic relationships of the Orchidaceae at mid and deep evolutionary time scales. More recently, phylogenies have frequently been employed as a source of information to conduct taxonomic transfers in the Orchidaceae. While this is a welcome line of evidence, extra caution must be exerted whenever such changes are proposed based on phylogenies containing significant taxon sampling gaps and insufficient statistical support (e.g., **Szlachetko et al., 2013; 2022**). Phylogenetic relationships can substantially change whenever extended taxon sampling and / or informative sequences are considered in new analyses (**Zwickl and Hillis 2002; Heath et al., 2008**). To avoid any unnecessary confusion in generic/subtribal/tribal taxonomic delimitation (e.g., **Szlachetko et al., 2013; 2022**) at present, we advise against any taxonomic transfers informed by our phylogeny as it currently lacks sufficient taxonomic representation (and in some cases support) at multiple phylogenetic levels.

***Stem and crown node absolute time discordance***

The comparison of stem ages derived from the Angiosperm353 chronograms and crown node ages of the Sanger chronogram indicated that age discordance occurred only in 8% of all Sanger genera grafted into the Angiosperms353 chronogram (**Figure S11**). This is a strong indication that time discordance is not frequent between the Angisoperm353 and Sanger datasets and, thus is unlikely to bias our tree-merging approach.

**References**

Balbuena JA, Miguez-Lozano R, Blasco-Costa I. 2013. PACo: a novel Procustres application to cophylogenetic analysis. *PLoS ONE* 8(4): e61408

Chase MW, Cameron KM, Freudenstein JV, Pridgeon AM, Salazar G, van den Berg C, Schuiteman A. 2015. An updated classification of Orchidaceae. *Botanical Journal of the Linnean Society* 177: 151-174.

Doyle JJ, Doyle JL. 1990. Isolation of plant DNA from fresh tissue. *Focus* 12: 13-15

Heath TA, Hedtke SM, Hillis D. 2008. Taxon sampling and the accuracy of phylogenetic analysis. *Journal of Systematics and Evolution* 46(3): 239-257.

Górniak M, Paun O, Chase MW. 2010. Phylogenetic relationships within Orchidaceae based on a low-copy nuclear coding gene, *Xdh*: congruence with organellar and nuclear ribosomal DNA results. *Molecular Phylogenetics and Evolution* 56(2): 784-795.

Johnson MG, Gardner EM, Liu Y, Medina R, Goffinet B, Shaw AJ, Zerega NJ, Wicket NJ. 2016. HybPiper: extracting coding sequence and introns for phylogenetics from high-throughput sequencing reads using target enrichment. *Applications in Plant Sciences* 4(7): 1600016

Kores PJ, Molvray M, Weston PH, Hopper SD, Brown AP, Cameron KM, Chase MW. 2001. A phylogenetic analysis of Diurideae (Orchidaceae) based on plastid DNA sequence data. *American Journal of Botany* 88(10): 1903-1914.

Li H, Dublin R. 2009. Fast and accurate short read alignment with Burrows-Wheeler transform. *Boinformatics* 25:1754-1760.

Li Y-X, Li Z-H, Schuiteman A, Chase MW, Li J-W, Huang W-Ch, Hidayat A, Wu Sh-Sh, Jin X-H. 2019. Phylogenomics of Orchidaceae based on plastid and mitochondrial genomes. *Molecular Phylogenetics and Evolution* 139: 106540

Mauad AVS, Vieira LN, Baura VA, Balsanelli E, de Souza EM, Chase MW, Smidt EC. 2021. *PLoS ONE* 16(8): e0256126

McLay TGB, Birch JL, Gunn BF, Ning W, Tate JA, Nauheimer L, Joyce EM, Simpson L, Schmidt-Lebuhn A, Baker WJ, Forest F, Jackson CJ. 2021. New target acquired: improving locus recovery from the Angiosperms353 probe set. *Applications in Plant Sciences* 9(7): 1-9

Paradis E, Claude J, Strimmer K. 2004. APE: analyses of phylogenetic and Evolution in R language. *Bioinformatics* 20: 289 – 290.

Pérez-Escobar OA, Balbuena JA, Gottschling M. 2016. Rumbling orchids: how to assess divergent evolution between chloroplast endosymbionts and the nuclear host. *Systematic Biology* 65(1): 51-65.

Pérez-Escobar OA, Gottschling M, Chomicki G, Condamine FL, Klitgaard B, Pansarin E, Gerlach G. 2017. Andean mountain building did not preclude dispersal of lowland epiphytic orchids in the Neotropics. *Scientific Reports* 7(1): 4919.

Pérez-Escobar OA, Dodsworth S, Bogarín D, Balbuena JA, Schley RJ, Kikuchi IZ, Morris SK, Epitawalage N, Cowan R, Maurin O, Zuntini A, Arias T, Serna-Sánchez A, Gravendeel B, Torres MF, Nargar K, Chomicki G, Chase MW, Leitch IJ, Forest F, Baker WJ. 2021a. Hundreds of nuclear and plastid loci yield novel insights into orchid relationships. *American Journal of Botany* 108(7): 1166-1180.

R Core Team (2022). R: a language and environment for statistical computing. R Foundation for Statistical Computing, Vienna, Austria.

Serna-Sánchez M, Pérez-Escobar OA, Bogarín D, Torres-Jimenez MF, Alvarez-Yela AC, Arcila-Galvis JE, Hall C, de Barros D, Pinheiro F, Dodsworth S, Chase MW, Antonelli A, Arias T. 2021. Plastid phylogenomics resolves ambiguous relationships within the orchid family and provides a solid timeframe for biogeography and macroevolution. *Scientific Reports* 11: 6858

Smith SA, Moore MJ, Brown JW, Yang Ya. 2015. Analysis of phylogenomic datasets reveal conflict, concordance, and gene duplications with examples from animals and plants. *BMC Evolutionary Biology* 15: 150

Szlachetko DL, Tukallo P, Mytnik-Ejsmont, Grochocka E. 2013. Reclassification of the *Angraecum* alliance (Orchidaceae, Vandoideae) based on molecular and morphological data. *Biodiversity Research* 29: 1-23.

Szlachetko DL, Dudek M, Naczk A, Kolanowska M. 2022. Taxonomy and biogeography of *Andinia* complex (Orchidaceae). *Diversity* 14(5): 372

Stull GW, Pham KK, Soltis PS, Soltis DE. 2023. Deep reticulation: the long legacy in vascular plant evolution. *The Plant Journal* doi.org/10.1111/tpj.16142.

Van der Niet T, Linder PH. 2008. Dealing with incongruence in the quest of the species tree: a case of study from the orchid genus Satyrium. *Molecular Phylogenetics and Evolution* 47: 154-174.

Vargas OM, Ortiz EM, Simpson BB. 2017. Conflicting phylogenomic signals reveal a pattern of reticulate evolution in a recent high-Andean diversification (Asteraceae: Astereae: *Diplostephium*). *New Phytologist* 214(4): 1736- 1750.

Zhang C, Rabiee M, Sayyari E, Mirarab S. 2018. ASTRAL-III: polynomial time species tree reconstruction from partially resolved gene trees. *BMC Bioinformatics* 19:153.

Zhang G, Hu Y, Huang M-Z, Huang W-C, Liu D-K, Zhang D, Hu H, Downing JL, Liu Z-J, Ma H. 2023. Comprehensive phylogenetic analyses of Orchidaceae using nuclear genes and evolutionary insights into epiphytism. *Journal of Integrative Plant Biology*, doi.org/10.1111/jipb.13462

Zwickl D, Hillis DM. 2002. Increased taxon sampling greatly reduces phylogenetic error. *Sytematic Biology* 51(4): 588-598.

**Supplementary Figures**

**Figure S1.** A detailed view of phylogenetic relationships between the Apostasioideae, Vanilloideae, and Cypripedioideae subfamilies. **A**) A zoom-in of the consensus tree-network inferred from 200 bootstrap replicate maximum likelihood (ML) trees derived from the concatenated alignment of 339 low-copy nuclear genes. Circles at nodes represent bipartitions present in > 75% of the bootstrap ML trees. Non-monophyletic groups are highlighted in bold and grey. Samples sequenced from typological material are highlighted in bold and pink. **B**) General view of the consensus tree-network. **C**) Multispecies-coalescent tree, computed from 339 nuclear ML gene trees in the software ASTRAL-III. Pie charts at nodes indicate the proportion of gene trees that support (blue), or are in conflict with (green, red) the species tree. Here, the green proportion of the pie chart indicates the proportion of gene trees recovering the second most frequent alternative bipartition to the species tree, and red – any other alternative conflicting bipartition. Grey represents non-informative gene trees. **D**) Proportion of missing sequence data per sample. Blue and yellow bars represent informative gene sequences and the proportion of missing sequences, respectively. Photos: Oscar A. Pérez-Escobar, Kerry Dressler & Diego Bogarín.

**Figure S2. A**) A detailed view of phylogenetic relationships within the Orchidoideae subfamily. **A**) A zoom-in of the consensus tree-network inferred from 200 bootstrap replicate maximum likelihood (ML) trees derived from the concatenated alignment of 339 low-copy nuclear genes. Circles at nodes represent bipartitions present in > 75% of the bootstrap ML trees. Non-monophyletic groups are highlighted in bold and grey. Samples sequenced from typological material are highlighted in bold and pink. **B**) General view of the consensus tree-network. **C**) Multispecies-coalescent tree, computed from 339 nuclear ML gene trees using the software ASTRAL-III. Pie charts at nodes indicate the proportion of gene trees that support (blue), or are in conflict with (green, red) the species tree. Here, the green proportion of the pie chart indicates the proportion of gene trees recovering the second most frequent alternative bipartition to the species tree, red – any other alternative conflicting bipartition. Grey represents non-informative gene trees. **D**) Proportion of missing sequence data per sample. Blue and yellow bars represent informative gene sequences and the proportion of missing sequences, respectively. Photos: Oscar A. Pérez-Escobar.

**Figure S3. A**) A detailed view of phylogenetic relationships between early diverging Epidendroideae lineages. **A**) A zoom-in of the consensus tree-network inferred from 200 bootstrap replicate maximum likelihood (ML) trees derived from the concatenated alignment of 339 low-copy nuclear genes. Circles at nodes represent bipartitions present in > 75% of the bootstrap ML trees. Non-monophyletic groups are highlighted in bold and grey. Samples sequenced from typological material are highlighted in bold and pink. **B**) General view of the consensus tree-network. **C**) Multispecies-coalescent tree, computed from 339 nuclear ML gene trees in the software ASTRAL-III. Pie charts at nodes indicate the proportion of gene trees that supports (blue), or conflicts (green, red) with the species tree. Here, the green proportion of the pie diagram indicates the proportion of gene trees recovering the second most frequent alternative bipartition to the species tree, and red – any other alternative conflicting bipartition. Grey represents non-informative gene trees. **D**) Proportion of missing sequence data per sample. Blue and yellow bars represent informative gene sequences and the proportion of missing sequences, respectively. Photos: Oscar A. Pérez-Escobar.

**Figure S4. A**) A detailed view of phylogenetic relationships within the Cymbidieae. **A**) A zoom-in of the consensus tree-network inferred from 200 bootstrap replicate maximum likelihood (ML) trees derived from the concatenated alignment of 339 low-copy nuclear genes. Circles at nodes represent bipartitions present in > 75% of the bootstrap ML trees. Non monophyletic groups are highlighted in bold and grey. Samples sequenced from typological material are highlighted in bold and pink. **B**) General view of the consensus tree-network. **C**) Multispecies-coalescent tree, computed from 339 nuclear ML gene trees in the software ASTRAL-III. Pie charts at nodes indicate the proportion of gene trees that support (blue), or are in conflict with (green, red) with the species tree. Here, the green proportion of the pie chart indicates the proportion of gene trees recovering the second most frequent alternative bipartition to the species tree, and red – any other alternative conflicting bipartition. Grey represents non-informative gene trees. **D**) Proportion of missing sequence data per sample. Blue and yellow bars represent informative gene sequences and the proportion of missing sequences, respectively. Photos: Oscar A. Pérez-Escobar.

**Figure S5. A**) A detailed view of phylogenetic relationships within the Vandeae. **A**) A zoom-in of the consensus tree-network inferred from 200 bootstrap replicate maximum likelihood (ML) trees derived from the concatenated alignment of 339 low-copy nuclear genes. Circles at nodes represent bipartitions present in > 75% of the bootstrap ML trees. Non monophyletic groups are highlighted in bold and grey. Samples sequenced from typological material are highlighted in bold and pink. **B**) General view of the consensus tree-network. **C**) Multispecies-coalescent tree, computed from 339 nuclear ML gene trees in the software ASTRAL-III. Pie charts at nodes indicate the proportion of gene trees that supports (blue), or in in conflict with (green, red) the species tree. Here, the green proportion of the pie charts indicates the proportion of gene trees recovering the second most frequent alternative bipartition to the species tree, and red – any other alternative conflicting bipartition. Grey represents non-informative gene trees. **D**) Proportion of sequence data missing per sample. Blue and yellow bars represent informative gene sequences and the proportion of missing sequences, respectively. Photos: Oscar A. Pérez-Escobar, Ori Fragman & Greg Allikas.

**Figure S6. A**) A detailed view of phylogenetic relationships within the Epidendreae. **A**) A zoom-in of the consensus tree-network inferred from 200 bootstrap replicate maximum likelihood (ML) trees derived from the concatenated alignment of 339 low-copy nuclear genes. Circles at nodes represent bipartitions present in > 75% of the bootstrap ML trees. Non monophyletic groups are highlighted in bold and grey. Samples sequenced from typological material are highlighted in bold and pink. **B**) General view of the consensus tree-network. **C**) Multispecies-coalescent tree, computed from 339 nuclear ML gene trees in the software ASTRAL-III. Pie charts at nodes indicate the proportion of gene trees that support (blue), or are in conflict with (green, red) the species tree. Here, the green proportion of the pie diagram indicates the proportion of gene trees recovering the second most frequent alternative bipartition to the species tree, and red – any other alternative conflicting bipartition. Grey represents non-informative gene trees. **D**) Proportion of missing sequence data per sample. Blue and yellow bars represent informative gene sequences and the proportion of missing sequences, respectively. Photos: Oscar A. Pérez-Escobar.

**Figure S7.** A genus-level chronogram of the Orchidaceae produced from 25 low copy nuclear, clock-like genes in 339 samples. Branch colour denotes branch support (posterior probability). (Inset:) 3D dotplot of posterior probability support along branches vs branch length (million years) and note height (million years ago [Ma]) as seen from two angles. The dotplot reveals that low posterior probability values are restricted to short branches diverging mostly 50 Ma to present.

**Figure S8.** Species-level Maximum Clade Credibility ultrametric tree of the orchid family, derived from the grafting of 500 MCMC trees produced from a *matK*-ITS supermatrix (1940 samples) onto 500 MCMC trees produced from 25 clock-like genes (339 samples). Branch colour denotes branch support (posterior probability). (Inset:) 3D dotplot of posterior probability support along branches vs. branch length (million years) and node height (million years ago [Ma]). The dotplot reveals that low posterior probability values are restricted to short branches diverging mostly 30 Ma ago to the present.

**Figure S9.** The age of modern orchid diversity (genera and species) as inferred from branch lengths obtained from the ten PP species trees.

**Figure S10.** Global patterns of species richness per botanical country calculated from the WCVP database; warm colours indicate higher numbers of species per botanical country, whereas cold colours indicate lower numbers of species. Grey colours represent botanical countries without data.

**Figure S11. A)** Genera crown and stem node age congruence in the Sanger species level and NGS genus level consensus phylogenies (as shown in Figure S7, S8). **B)** Frequency of stem and crown node ages obtained respectively from the ten randomly sampled NGS genus level and Sanger species level posterior probability trees.

**Supplementary Tables**

**Table S1.** Voucher information, taxonomic rank, and proportion of informative and missing Angiosperm353 sequences of the plant material included in the construction of the NGS phylogenomic backbone. Due to the size of this data frame, the table is available at10.6084/m9.figshare.22245940.

**Table S2.** Genbank accession numbers of the samples mined from Genbank and retained for subsequent analysis. The sequences included in the species-level phylogeny are provided in Table S7. Due to the size of this data frame, the table is available at10.6084/m9.figshare.22245940.

**Table S3.** Root to tree variance, tree length, and bipartition support of the Angiosperm353 genes as inferred by SortAdate. Due to the size of this data frame, the table is available at10.6084/m9.figshare.22245940.

**Table S4.** The number of informative and missing sequences included in the inference of absolute age estimation analyses for the genus-level orchid backbone. Due to the size of this data frame, the table is available at 10.6084/m9.figshare.22245940.

**Table S5.** Ancestral areas and their corresponding probabilities of key nodes of the orchid species PP trees, as inferred by DEC. Detailed results for each ancestral area estimation are available at at10.6084/m9.figshare.22245940. Due to the size of this data frame, the table is available at10.6084/m9.figshare.22245940.

**Table S6.** Orchid species richness per botanical country as calculated from the WCVP. at10.6084/m9.figshare.22245940

**Table S7.** Maximum, minimum, and mean tip speciation rates derived from the ten PP species trees using the software BAMM. at10.6084/m9.figshare.22245940

**Supplementary Data S1**

Angiosperm353 alignments, sampling fractions per genera employed in BAMM, the ten PP species trees and their corresponding ancestral area estimations and speciation rate analyses are provided in Figshare link at10.6084/m9.figshare.22245940
